# The G protein-coupled receptor TBXA2R activates ERMs to control cell motility and invasion of triple-negative breast cancer cells

**DOI:** 10.1101/2023.03.28.534587

**Authors:** Kévin Leguay, Omaima Naffati, Yu Yan He, Mireille Hogue, Chloé Tesnière, Melania Gombos, Hellen Kuasne, Louis Gaboury, Christian Le Gouill, Sylvain Meloche, Michel Bouvier, Sébastien Carréno

**Author notes:** Corresponding authors: M. Bouvier, S. Carréno. These authors contributed equally to this work (Michel Bouvier, Sébastien Carréno).

## Abstract

Cell migration and invasion are critical processes for cancer cell metastasis, relying on the ability of cells to adapt their morphology. Proteins of the ezrin, radixin, and moesin (ERM) family are key regulators of cell morphogenesis and essential determinants of cancer cell metastasis. However, the mechanisms by which ERMs are activated in metastatic cells remain poorly understood. Here, we identify the thromboxane A2 receptor (TBXA2R), a G protein-coupled receptor overexpressed in multiple cancers, as a critical activator of ERMs, enhancing the motility and invasion of triple-negative breast cancer (TNBC) cells. We found that TBXA2R activates ERMs by engaging the Gα_q/11_ and Gα_12/13_ subfamilies, the small GTPase RhoA, and its Ser/Thr kinase effectors SLK and LOK. Furthermore, we demonstrate that TBXA2R promotes TNBC cell motility and invasion *in vitro* and metastatic colonization *in vivo*, dependent on ERM function. These findings reveal a novel signaling axis by which a member of the largest class of receptors activates key metastatic determinants, thereby controlling various aspects of metastasis. This discovery opens new avenues for developing targeted therapies against cancer metastasis.

## INTRODUCTION

Ezrin, radixin, and moesin (ERMs) are a family of membrane-cytoskeleton linker proteins that regulate cell morphogenesis, migration, and invasion (Fehon et al., 2010). They integrate actomyosin forces and microtubule signaling at the cell cortex by crosslinking actin filaments (Algrain et al., 1993) and microtubules (Solinet et al., 2013) with the plasma membrane. Inactive ERMs cycle between the cytoplasm and the plasma membrane in a closed conformation, where their N-terminal FERM domain binds their C-terminal C-ERMAD domain (Leguay et al., 2021). Upon stimulation, ERMs open up and expose their FERM microtubule-binding domain (Solinet et al., 2013) and their C-ERMAD actin-binding site (Gary and Bretscher, 1995). The C-ERMAD also harbors a conserved regulatory threonine residue (T567, T564, and T558 in ezrin, radixin, and moesin, respectively) that is phosphorylated upon stimulation to maintain the active open conformation (Simons et al., 1998). ERMs were shown to control cell morphogenesis during mitosis (Carreno et al., 2008; De Jamblinne et al., 2020; Kunda et al., 2008; Leguay et al., 2022; Roubinet et al., 2011), cell migration (Arpin et al., 2011; Barik et al., 2022; Hoskin et al., 2015) and cell invasion (Estecha et al., 2009; Song et al., 2020). In a pathological context, high levels of ezrin, radixin, or moesin expression correlate with a high rate of metastasis and poor prognosis in several cancers (Clucas and Valderrama, 2014). For instance, increased expression of moesin correlates with metastatic progression of oral squamous cell carcinoma (Kobayashi et al., 2004), melanoma (Estecha et al., 2009), and breast carcinoma (Bartova et al., 2017). Radixin is overexpressed in prostate cancer (Bartholow et al., 2011). Ezrin is overexpressed and hyperactivated in several cancer cells, including metastatic rhabdomyosarcoma (Yu et al., 2004), osteosarcoma (Khanna et al., 2004), prostate cancer (Pang et al., 2004) and mammary carcinoma (Bruce et al., 2007). In mice, ERM inhibition prevents breast cancer or osteosarcoma metastasis (Elliott et al., 2005; Ghaffari et al., 2019). These examples and other observations led to the notion that ERMs play central roles in metastasis (Barik et al., 2022; Curto and McClatchey, 2004). ERMs are, therefore, attractive targets for anti-cancer therapies. However, the mechanisms by which ERMs become activated during metastasis remain unknown.

Metastasis is responsible for most cancer deaths (Seyfried and Huysentruyt, 2013). This complex process is driven by the dysregulation of cell motility, which usually plays a crucial role in development and tissue repair. In cancer cells, aberrant signaling pathways activate cytoskeletal rearrangements to promote cell migration and enable invasion and dissemination to distant sites (Fife et al., 2014). In the last decade, evidence showed that cancer cells do not act alone but dialogue with surrounding cells and extracellular matrix components that promote tumor progression and metastasis (Clark and Vignjevic, 2015). The thromboxane A2 receptor (TBXA2R) is a G protein-coupled receptor (GPCR) that mediates platelet activation and aggregation in response to thromboxane A2 (TXA2), a short half-life (∼30 s) prostaglandin derivative (Jones et al., 1985). TXA2 also controls other functions, such as vasoconstriction, inflammation, and angiogenesis (Nakahata, 2008). In a pathological context, TXA2 promotes metastasis of several cancers, including colorectal (Guillem-Llobat et al., 2016), lung (Lucotti et al., 2019), prostate (Nie et al., 2004) and breast (Ekambaram et al., 2011) cancer. In addition, TBXA2R overexpression correlates with poor prognosis in aggressive breast tumors (Watkins et al., 2005), and targeting the TXA2 pathway has been proposed as a strategy against metastasis of these tumors (Li et al., 2017).

Breast cancer is a very heterogeneous disease that can be classified into different subtypes based on the expression levels of different receptors. Among these subtypes, triple-negative breast cancer (TNBC) does not express estrogen, progesterone, and HER2 receptors but do overexpress TBXA2R (Orr et al., 2016). Both the high metastatic properties of TNBC cells and the lack of efficient, targeted therapy explain the high mortality rate of this cancer. While TNBC represents 10 to 20% of new breast cancer cases, it represents more than 30% of death associated with the disease (Bianchini et al., 2022). The molecular mechanisms underlying TNBC metastasis is still poorly understood; thus, identifying the mechanisms that prompt these cells to disseminate could lead to developing more effective treatments for TNBC patients.

We recently demonstrated that RhoA, a small GTPase, activates ERMs at the onset of mitosis (Leguay et al., 2022). Interestingly, several GPCRs, including TBXA2R, are known to engage RhoA to execute some of their functions (Yu and Brown, 2015). This prompted us to investigate the potential functional relationship between TBXA2R, RhoA, and ERMs. We discovered that TBXA2R engages the heterotrimeric G proteins Gα_q/11_ and Gα_12/13_ subfamilies, RhoA, and its kinase effectors SLK and LOK to activate ERMs in TNBC cells. Furthermore, we showed that activation of TBXA2R in these cells enhances cell migration and invasion *in vitro* and metastatic colonization in a mouse model of metastasis by a mechanism dependent on the activation of ERMs.

## RESULTS

### Activation of TBXA2R promotes ERM opening and activation

To determine if TBXA2R activates ERM proteins, we took advantage of ERM conformational enhanced bystander BRET (ebBRET) biosensors that monitor the opening of individual ERMs at the plasma membrane (Leguay et al., 2021). The C-terminus of ezrin, radixin or moesin is fused to the bioluminescent energy donor *Renilla* luciferase (rLucII) while their N-terminus is anchored at the plasma membrane using a myristoylation signal and polybasic motif. The fluorescent energy acceptor *Renilla* GFP (rGFP) is also targeted to the plasma membrane through its fusion with the prenylated CAAX motif of KRAS (rGFP-CAAX) (Fig S1A). In their closed, inactive conformation, ERMs are globular with a diameter of ∼10 nm, while they open up to ∼40 nm when activated (Liu et al., 2007). Since the transfer of energy between the BRET donor and acceptor is inversely proportional to the distance separating them (Breton et al., 2010), the opening and activation of ERMs result in a decrease in ebBRET signals (Leguay et al., 2021). We first assessed the ability of TBXA2R to promote ERM opening and activation in HEK293T cells, as these cells are well suited to study GPCR signaling using BRET assays (Breton et al., 2010; Gales et al., 2005; Kobayashi et al., 2019; Namkung et al., 2016; Namkung et al., 2018). Since TBXA2R is not expressed in these cells (Atwood et al., 2011), we co-transfected this GPCR with individual ERM biosensors and measured their ebBRET signal. This approach revealed that the three ERM proteins are opened downstream of TBXA2R. Activating this GPCR using U46619, a stable TBXA2R agonist (Coleman et al., 1981), decreased ebBRET signals associated with each ERM biosensor (Fig 1A and S1B-C). U46619 promoted ezrin, radixin, or moesin opening with similar low nanomolar potency (Fig 1B), in agreement with the affinity of this agonist for TBXA2R (Kattelman et al., 1986). The phosphorylation of a conserved threonine residue stabilizes the ERM open, active form. We assessed this last step of ERM activation using an antibody that specifically recognizes phosphorylated ERM (p-ERM) (Carreno et al., 2008). Confirming the results obtained with the ERM biosensors, U46619 promoted a substantial increase in phosphorylation of endogenous ERMs, comparable to the increase triggered by calyculin A, a potent inhibitor of Ser/Thr phosphatases (Fig 1C-D). Furthermore, we found that ERM activation downstream of TBXA2R depends on the catalytic activity of a Ser/Thr kinase since staurosporine, a broad-spectrum kinase inhibitor prevents ERM phosphorylation upon U46619 treatment (Fig 1C-D). Consistent with these findings, staurosporine also prevented the U46619-induced ebBRET decrease in agreement with the role of phosphorylation in stabilizing the ERM open conformation (Fig 1A and S1B-C). We then characterized the kinetics of ERM activation downstream of TBXA2R and showed that U46619 triggers ERM phosphorylation very rapidly as the levels of p-ERM reach their maximum at 2 minutes, the earliest time-point tested. We also showed that this activation lasts at least 4 hours (Fig 1E-F).

**FIGURE 1:**
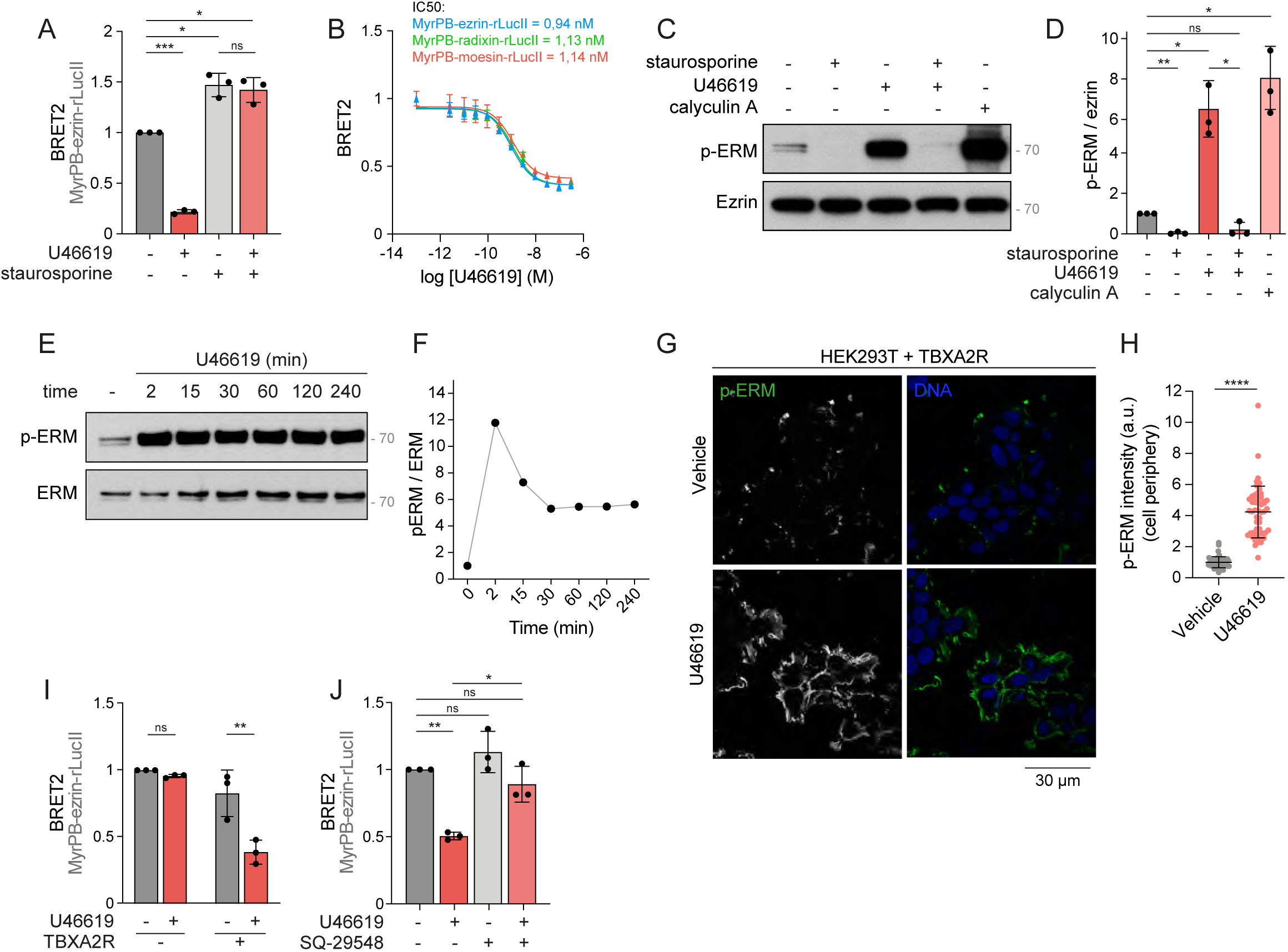
TBXA2R stimulation triggers ERM activation in HEK293T cells. (A) ebBRET signals were measured in HEK293T cells expressing MyrPB-ezrin-rLucII, co- transfected with TBXA2R (HEK293T-TBXA2R) and treated with vehicle, 100 nM U46619 for 5 min and/or 100 nM staurosporine for 30 min. (B) ebBRET signals measured in HEK293T-TBXA2R cells expressing MyrPB-E,R,M-rLucII and treated with increasing concentrations of U46619 for 5 min. (**C-D**) Immunoblot of HEK293T-TBXA2R cells treated with vehicle, 100 nM staurosporine for 30 min, 100 nM U46619 for 5 min, and/or 100 nM calyculin A for 10 min (**C**). p-ERM over ezrin signals were quantified and normalized to the vehicle (**D**). (**E-F**) Immunoblot of HEK293T-TBXA2R cells treated with 100 nM U46619 for the indicated times (**E**). p-ERM over ERM signals were quantified and normalized to the vehicle (**F**). (**G-H**) Immunofluorescence of HEK293T-TBXA2R cells treated with vehicle or 100 nM U46619 for 5 min (**G**). p-ERM staining at the cell periphery was quantified and normalized to cells treated with vehicle (**H**). (I) ebBRET signals measured in HEK293T cells expressing MyrPB-ezrin-rLucII and co- transfected with or without TBXA2R and treated with vehicle or 100 nM U46619 for 5 min. (J) ebBRET signals were measured in HEK293T-TBXA2R cells expressing MyrPB-ezrin-rLucII biosensor and treated with vehicle, 100 nM U46619 for 5 min and/or 1 µM SQ-29548 for 30 min. *ebBRET signals (**A-B, I-J**) represent the mean +/- s.d. of three independent experiments. Immunoblots (**C, E**) and immunofluorescences (**G**) are representative of three independent experiments. P-ERM quantifications (**D, H**) represent the mean +/- s.d. of three independent experiments. P-ERM quantification (**F**) is representative of three independent experiments. Dots represent independent experiments (**A, D, I-J**), individual cells (**H**), or the mean of independent experiments (**B, F**). P values were calculated using one-sample t-test (**J**), two-tailed unpaired t-test (**H**), or two-tailed paired t-test (**A, D, I**) except for comparison made with normalizing condition (vehicle) where one-sample t-test was applied. *, P < 0.05. **, P < 0.01. ***, P < 0.001. ****, P < 0.0001. ns, not significant*.

Finally, immunofluorescence analysis of p-ERM revealed that TBXA2R activation increases ERM phosphorylation at the cell cortex, where ERMs crosslink the cytoskeleton with the plasma membrane (Fig 1G-H). Altogether, these experiments show that activation of TBXA2R promotes the opening and sustained activation of ERMs.

### TBXA2R engages Gα_q/11_, Gα_12/13_, RhoA, and SLK kinases to activate ERMs

Taking advantage of the versatility of HEK293 cells for transfection and loss-of-function experiments (Pulix et al., 2021), we aimed to identify the signaling pathway that TBXA2R engages in activating ERMs. Since TBXA2R activates ezrin, radixin, and moesin similarly (Fig 1B), we focused on the ezrin biosensor as a proxy to study ERM regulation. In accordance with U46619 activating ERMs through TBXA2R, U46619 did not affect ebBRET signals associated with the ezrin biosensor in cells that were not transfected with TBXA2R (Fig 1I). Further demonstrating the specificity of this newly discovered signaling axis, SQ- 29548, a highly selective TBXA2R antagonist (Ogletree et al., 1985), prevented ezrin opening upon TBXA2R activation with U46619 (Fig 1J).

When activated, TBXA2R engages Gα_q/11_ and Gα_12/13_ sub-family members that relay the signal to different effectors (Avet et al., 2020; Nakahata, 2008). To characterize the role of each of these two Gα sub-families in ERM activation, we used CRISPR-engineered HEK293 cells lacking the Gα_q/11_ (Schrage et al., 2015) or Gα_12/13_ (Devost et al., 2017) subunits (ΔGα_q/11_ and ΔGα_12/13_ cells, respectively). In ΔGα_q/11_ cells, TBXA2R activation still promoted full ezrin opening while deletion of Gα_12/13_ subunits partially inhibited this opening (Fig 2A). We then tested whether Gα_q/11_ and Gα_12/13_ sub-families exert redundant functions in ERM activation downstream of TBXA2R. Treatment of ΔGα_12/13_ cells with YM- 254890, a selective Gα_q/11_ subunits inhibitor (Takasaki et al., 2004), completely inhibited ezrin opening upon TBXA2R activation (Fig 2A). This Gα_q/11_ inhibitor did not affect ezrin opening in control parental cells downstream of TBXA2R activation, consistent with the results obtained with the deletion of Gα_q/11_ subunits and confirming that either Gα_q/11_ or Gα_12/13_ subfamilies are sufficient to promote ERM opening.

**FIGURE 2:**
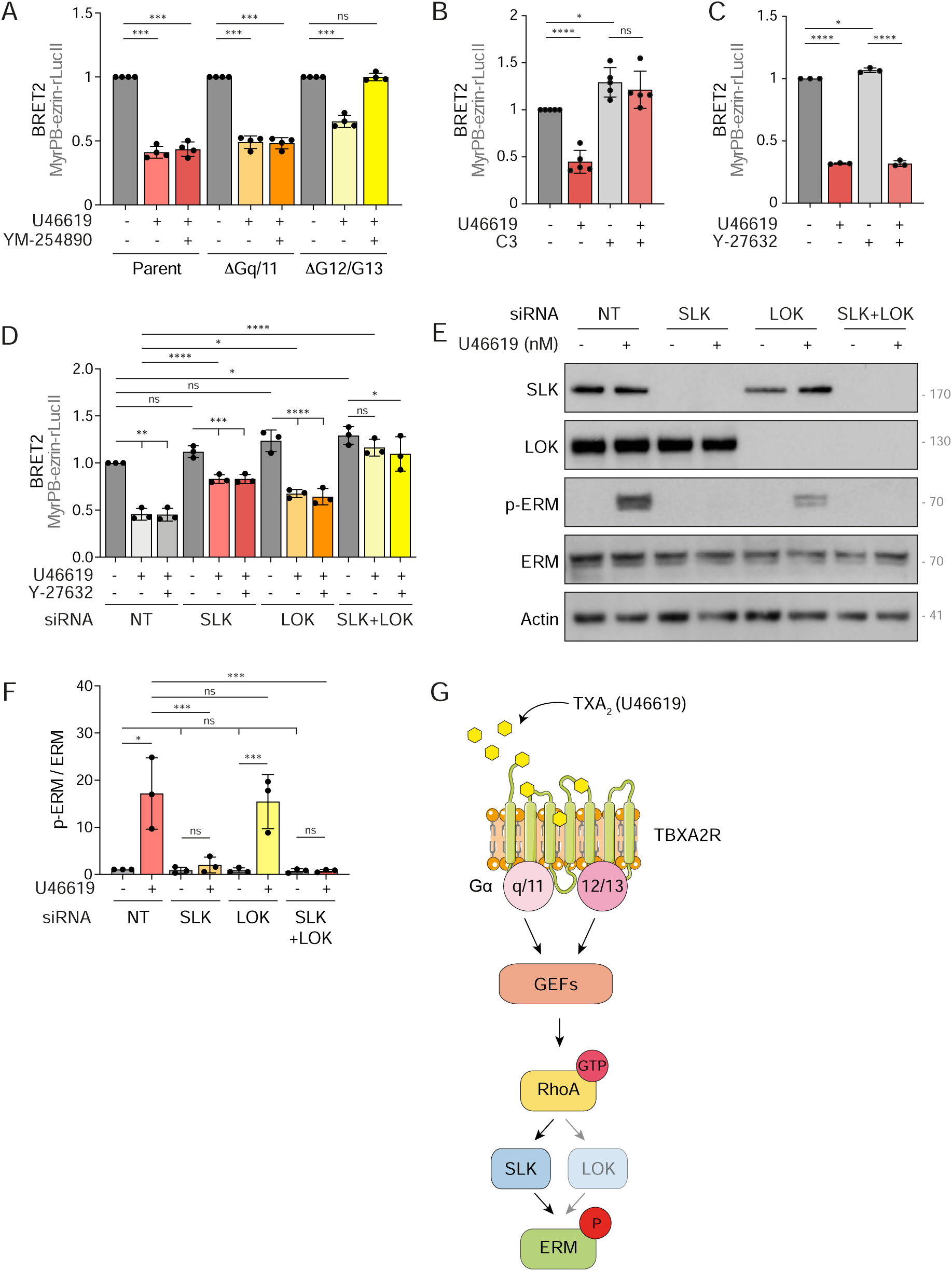
TBXA2R activates ERMs through RhoA and its kinase effectors SLK/LOK. **(A**) ebBRET signals measured in parental HEK293T cells or HEK293T cells knock-out for Gq/11 (ΔGq/11) or G12/13 (ΔG12/13), co-expressing TBXA2R and MyrPB-ezrin-rLucII biosensor and treated with 100 nM U46619 for 5 min and/or 1 µM YM-254890 for 30 min. ebBRET signals measured in each cell line are normalized to the vehicle in the same cell line. (**B-C**) ebBRET signals measured in HEK293T-TBXA2R expressing MyrPB-ezrin-rLucII biosensor and treated with 100 nM U46619 for 5 min and/or 1 µg/mL C3 transferase for 6h (**B**) or 10 µM Y-27632 for 4h (**C**). (**D**) ebBRET signals measured in HEK293T-TBXA2R expressing MyrPB-ezrin-rLucII biosensor, transiently transfected with non-target siRNA (NT) or siRNA targeting SLK and/or LOK and treated with 100 nM U46619 for 5 min and/or 10 µM Y-27632 for 4h. (**E-F**) Immunoblot of HEK293T-TBXA2R cells transiently transfected with non-target siRNA (NT) or siRNA targeting SLK and/or LOK and treated with 100 nM U46619 for 5 min (**E**). p- ERM over ERM signals were quantified and normalized to NT incubated with vehicle (**F**). (G) Proposed model for the signaling pathway downstream of TBXA2R that activates ERMs in HEK293T-TBXA2R cells. *ebBRET signals (**A-D**) represent the mean +/- s.d. of at least three independent experiments. Immunoblot (**E**) is representative of three independent experiments. P-ERM quantifications (**F**) represent the mean +/- s.d. of three independent experiments. Dots represent independent experiments (**A-D, F**). P values were calculated using a one-sample t-test (**A**), two-tailed paired t-test (**B-C**) or using Holm-Sidak’s multiple comparisons test with a single pooled variance (**D, F**) except for comparison made with normalizing condition (vehicle, **B**-**C**; NT + vehicle, **D,F**) where a one-sample t-test was applied. *, P < 0.05. **, P < 0.01. ***, P < 0.001. ****, P < 0.0001. ns, not significant*.

Among the effectors of Gα_q/11_ and Gα_12/13_, RhoA was shown to promote phosphorylation and activation of ERMs (Bagci et al., 2020; Leguay et al., 2022; Shaw et al., 1998). Indeed, treatment with the exoenzyme C3 transferase, a selective inhibitor of RhoA (Wilde and Aktories, 2001), prevented ezrin opening upon TBXA2R stimulation (Fig 2B), establishing that this small GTPase acts downstream of Gα_q/11_ and Gα_12/13_ subunits to activate ERMs. We next aimed to identifying which kinase(s) acts downstream of TBXA2R to activate ERMs. RhoA was shown to directly bind and activate two families of Ser/Thr kinases that can phosphorylate ERMs: the Rho-associated protein kinases ROCK1 and ROCK2 (Matsui et al., 1996) and the Ste20-like-kinase (SLK) and Lymphocyte-oriented-kinase (LOK) paralogs (Bagci et al., 2020; Leguay et al., 2022; Viswanatha et al., 2012). We previously showed that Y-27632, a specific ROCK inhibitor, inhibits ROCK kinases in HEK293T cells (Leguay et al., 2022). Here, we established that ROCK1 and ROCK2 activity is not essential to activate ERMs downstream of TBXA2R since Y-27632 did not affect ezrin opening following U46619 treatment (Fig 2C). In contrast, transient depletion of SLK by RNAi reduced ezrin opening downstream of TBXA2R activation (Fig 2D). While the depletion of LOK led to a slight inhibition of ezrin opening, the co-depletion of both LOK and SLK paralogs completely prevented ezrin opening downstream of TBXA2R (Fig 2D). Consistent with kinases of the SLK family being the sole kinases that relay the signal between TBXA2R and ERMs, inhibition of ROCK kinases with Y-27632 did not further reduce ezrin opening upon SLK, LOK or SLK/LOK transient depletion (Fig 2D). We then observed that LOK depletion has a minimal effect on ERM phosphorylation downstream of TBXA2R activation, as observed with the ezrin biosensor (Fig 2E-F). Conversely, SLK depletion almost completely abolished ERM phosphorylation, confirming the involvement of SLK as the primary kinase responsible for mediating the signal between TBXA2R and ERM activation in these cells. Furthermore, simultaneous depletion of SLK and LOK prevented any ERM phosphorylation, indicating that no other kinases are involved in this activation pathway.

Altogether, these results establish that TBXA2R activates RhoA through both Gα_q/11_ and Gα_12/13_, leading to ERM activation via kinases of the SLK family (Fig 2G).

### TBXA2R activates ERMs in TNBC cells

Having shown that TBXA2R activates ERMs, we hypothesized that this signaling axis could play a role in the biology of cancer cells. Given that TBXA2R and ERMs have previously been shown to regulate motility and invasion of TNBC cells in separate studies (Orr et al., 2016; Qin et al., 2020), we investigated the potential involvement of the TBXA2R-ERM signaling axis in human TNBC. Upon interrogation of transcriptomic data from TNBC samples obtained from the Cancer Genome Atlas of Invasive Breast Carcinoma (Cancer Genome Atlas, 2012), we observed that TNBC exhibited higher levels of TBXA2R mRNA expression compared to normal breast tissues (Fig 3A). Further analysis revealed that only a subset of TNBC patients exhibited higher mRNA expression levels of this GPCR compared to normal breast tissues (Fig 3B). Interestingly, our analyses showed that TBXA2R high mRNA expression is associated with poorer overall survival. Such association was not observed in non-TNBC subtypes (Fig 3C-D). To investigate whether ERMs are activated downstream of TBXA2R signaling in cancer tissues from TNBC patients, we examined the levels of phosphorylated ERMs in TNBC biopsies using tissue microarrays (TMA). Our analysis revealed a positive correlation between TBXA2R protein expression and levels of activated p-ERM in 55 TNBC samples (Fig 3E-F, r=0.50, p<0.0001). Based on the levels of p-ERM and TBXA2R protein expression, we identified three distinct subgroups of TNBC.

**FIGURE 3:**
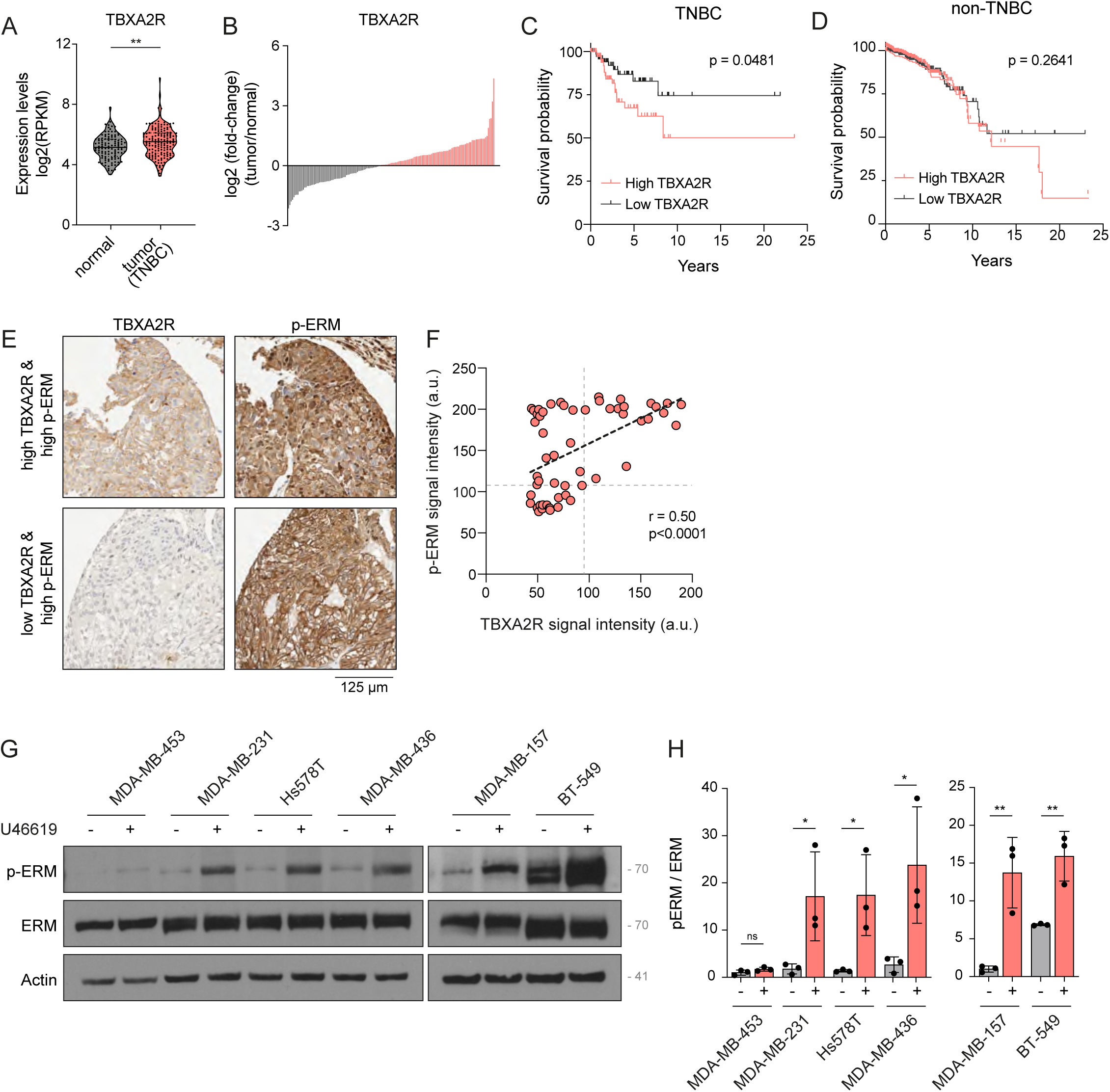
TBXA2R activates ERMs in TNBC cells. (**A-D**) Analysis of TBXA2R mRNA expression in TNBC cancer tissues and associated overall survival. Data were extracted from The Cancer Genome Atlas (TCGA). (**A**) TNBC breast cancers present higher TBXA2R mRNA expression when compared to normal breast tissue. (**B**) Within TNBC cases, a subset of tumors shows elevated TBXA2R mRNA expression relative to normal breast tissue. (**C**) Kaplan–Meier curves show that high expression of TBXA2R is associated with shorter overall survival in TNBC patients but not in non-TNBC patients. (**E-F**) Representative IHC staining of p-ERM and TBXA2R in triple-negative breast cancer tissue (**E**). p-ERM and TBXA2R signal intensities were quantified at the cell cortex (n=55, **F**). Linear regression and Pearson correlation score are represented on the graph. (**G-H**) Immunoblot of 6 TNBC cell lines treated with vehicle or 100 nM U46619 for 5 min (**G**). p-ERM over ERM signals were quantified and normalized to MDA-MB-453 treated with vehicle (left) or MDA-MB-157 treated with vehicle (right) (**H**). *Immunoblot (**G**) is representative of three independent experiments. P-ERM quantification* (H) *represents the mean +/- s.d. of three independent experiments. Dots represent independent samples (**A, F**) or individual experiment (**H**). P values were calculated using two-tailed unpaired t-test (**A**), Log-rank (Mantel-Cox) test (**C-D**), Pearson correlation (**F**), or two-tailed paired t-test (**H**). *, P < 0.05. **, P < 0.01. ns, not significant*.

The first subgroup exhibited low levels of both p-ERMs and TBXA2R, suggesting that ERM activation or TBXA2R plays a minor role, if any, in TNBC biology for this subgroup. The second showed high levels of p-ERMs but low TBXA2R expression, indicating possible alternative pathways for ERM activation. The third demonstrated high levels of both p- ERMs and TBXA2R, indicating that TBXA2R may activate ERMs in this subgroup. Notably, no TNBC samples exhibited high TBXA2R expression and low levels of p-ERMs, further supporting a role for TBXA2R signaling in ERM activation in TNBC.

Finally, we investigated whether TBXA2R promotes the phosphorylation of ERMs across a panel of different TNBC cell lines. We selected six TNBC cells lines based on the level of TBXA2R mRNA expression as referenced in the Cancer Dependency Map Portal (Tsherniak et al., 2017) (Fig S2A). Interestingly, higher expression levels of TBXA2R mRNA were associated with a greater ability of the U46619 agonist to promote ERM phosphorylation (Fig 3G-H). Consistent with the expected role of ERMs at the cell cortex, activation of endogenous TBXA2R by its agonist increased ERM phosphorylation at the plasma membrane of all examined TNBC cell lines (Fig S2B-M).

### TBXA2R activates ERMs through G_q/11_ and G_12/13_, RhoA and SLK/LOK in Hs578T TNBC cells

To further investigate the significance of the TBXA2R-ERM signaling axis in TNBC, we focused on the Hs578T cell line, which showed a median level of TBXA2R mRNA expression among the six TNBC cell lines tested (Fig S2A). Hs578T cells are well-characterized and have been extensively used in cancer research, providing a reliable model for studying the molecular mechanisms underlying cell motility, invasion and metastasis (Koedoot et al., 2019; Yankaskas et al., 2019). We first confirmed the pharmacological selectivity of the TBXA2R agonist in Hs578T cells by showing that the TBXA2R antagonist SQ29548 blocks U46619-promoted ERM phosphorylation downstream of TBXA2R(Fig 4A-B). As we observed in HEK293T cells, U46619 activated ERMs in Hs578T cells at nanomolar concentrations (Fig 1B and 4C-D). We tested the role of the two Gα sub-families using YM-254860 to inhibit Gα_q/11_ subunits and by overexpressing p115-RGS-CAAX, a previously characterized dominant-negative construct that inhibits Gα_12/13_ activity (Lukasheva et al., 2020). In this construct, the RGS domain of p115-RhoGEF that inactivates Gα_12/13_ signaling (p115-RGS) is targeted at the plasma membrane using a CAAX motif. As observed in HEK293T cells, inhibition of Gα_q/11_ alone, using YM254890, did not reduce the activation of ERMs downstream of TBXA2R. In contrast, inhibition of Gα_12/13_ partially prevented ERM activation as measured by western blot analysis of p-ERM (Fig 4E-F). Co-inhibition of both Gα sub-families further reduced ERM activation (Fig 4E-F). In addition, inhibition of RhoA using exoenzyme C3 transferase but not inhibition of ROCK kinases using Y-27632 prevented phosphorylation of ERMs upon TBXA2R activation (Fig 4G-H). Finally, confirming our findings in HEK293T cells, single CRISPR-Cas9 knockout of SLK or LOK was not sufficient to eliminate ERM phosphorylation downstream of TBXA2R activation (Fig 4I-J). However, the double knockout of the two Ser/Thr kinase orthologues completely abrogated ERM activation upon TBXA2R stimulation. Confirming the role of these kinases in the TBXA2R-RhoA signaling pathway, SLK kinase activity increased when the GPCR was activated by U46619 (Fig 4K). Taken together, these data establish that Gα_q/11_ and Gα_12/13_ subunits, RhoA, SLK and LOK, act downstream of TBXA2R to activate ERMs in Hs578T cells.

**FIGURE 4:**
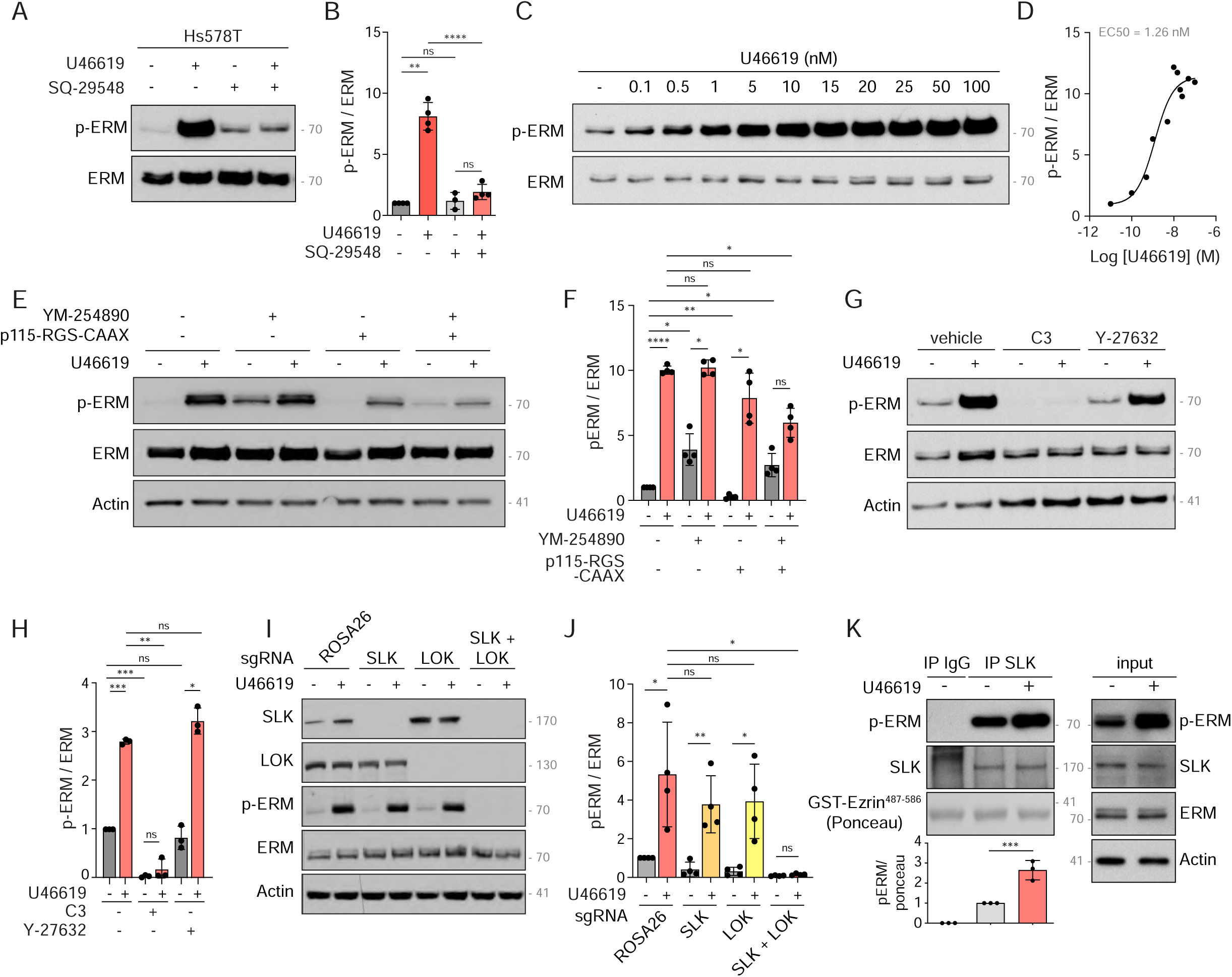
TBXA2R activates ERMs through Gα_q/11_ and Gα_12/13_, RhoA and SLK/LOK in Hs578T. (**A-B**) Immunoblot of Hs578T cells were treated with vehicle, 100 nM U46619 for 5 min and/or 1 µM SQ-29548 for 30 min (**A**). p-ERM over ERM signals were quantified and normalized to the vehicle (**B**). (**C-D**) Immunoblot of Hs578T cells treated with U46619 for 5 min with the indicated concentrations (**C**). p-ERM over ERM signals were quantified and normalized to the vehicle (**D**). (**E-H**) Immunoblots of Hs578T cells treated with vehicle or 100 nM U46619 for 5 min, 1 µM YM-254890 for 30 min, and/or transfected with 1 µg p115-RGS-CAAX (**E**), treated with vehicle or 100 nM U46619 for 5 min, 1 µg/mL C3 transferase for 6h and/or 10 µM Y-27632 for 4h (**G**). P-ERM over ERM signals were quantified and normalized to the vehicle (**F, H**). (**I-J**) Immunoblot of CRISPR-Cas9-mediated SLK and/or LOK Hs578T knockout cells treated with vehicle or 100 nM U46619 for 5 min (**I**). P-ERM over ERM signals were quantified and normalized to control cells (ROSA26) treated with vehicle (**J**). (K) Immune complex kinase assay of endogenous SLK immunoprecipitated from Hs578T cells incubated with vehicle or 100 nM U46619 for 5 min. The total lysate (input) is shown in the right panel. p-ERM signals over ponceau staining were quantified and normalized to immunoprecipitated SLK treated with vehicle (lane 2). *Immunoblots (**A, C, E, G, I, K**) are representative of at least two independent experiments. P-ERM quantifications (**B**, **D**, **F**, **H**, **J**, **K**) represent the mean +/- s.d. of at least two independent experiments. Dots represent independent experiments (**B, F, H, J, K**) or the mean of independent experiments (**D**). P values were calculated using Holm-Sidak’s multiple comparisons test with a single pooled variance (**B, F, H, J**) or a one-sample t-test (**K**) except for comparison made with normalizing condition (vehicle, **B, F, H**; sgROSA26 + vehicle, **J**) where one-sample t-test was applied. *, P < 0.05. **, P < 0.01. ***, P < 0.001. ****, P < 0.0001. ns, not significant*.

### TBXA2R activates SLK/LOK and moesin to potentiate Hs578T cell 2D motility, 3D invasion and metastatic colonization

We then investigated the role of the TBXA2R-ERM signaling axis in TNBC cell motility and invasion *in vitro*. To this end, we deleted individual ERMs by CRISPR-Cas9 editing of Hs578T cells. While the deletion of ezrin or radixin did not affect the phosphorylation of endogenous ERMs promoted by TBXA2R activation, the deletion of moesin almost totally abolished ERM phosphorylation (Fig 5A-B). We confirmed this finding by stably knocking-down ezrin, radixin, or moesin in Hs578T cells with shRNAs targeting each ERM (Fig S3A). Since TBXA2R activates ezrin, radixin, or moesin similarly in an engineered system(Fig 1B), this suggests that moesin is the most abundant ERM expressed in Hs578T cells. This hypothesis was confirmed by analyzing gene expression in Hs578T cells. Single-cell RNA sequencing data available in the Single Cell Expression Atlas (Papatheodorou et al., 2020)reveals that moesin mRNA is expressed at the highest level among the three ERMs in Hs578T cells (Fig S3B). We thus decided to further study the role of the TBXA2R-ERM signaling axis by using the Δmoesin Hs578T cells.

**FIGURE 5:**
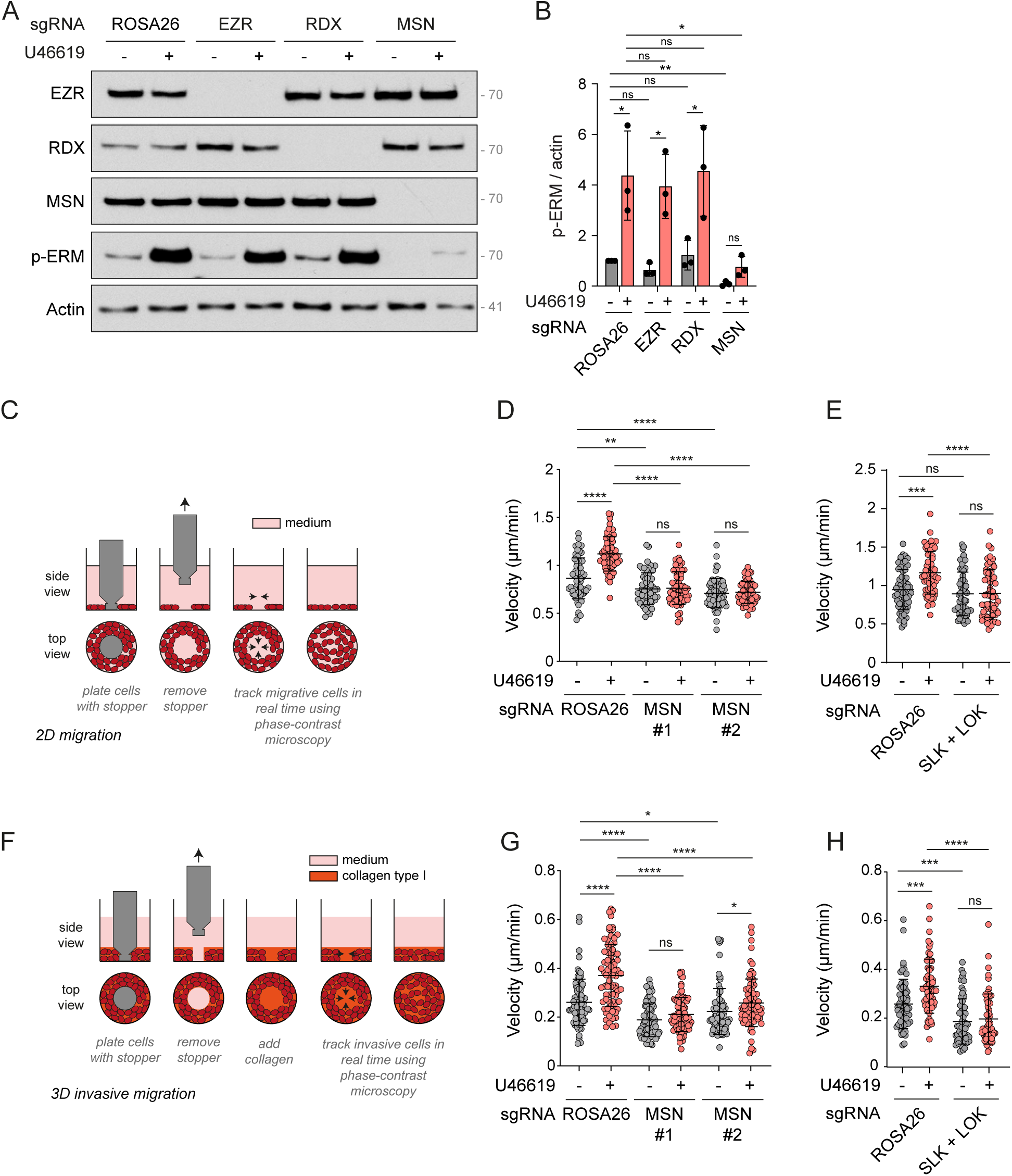
Moesin mediates TBXA2R-induced motility and invasion of Hs578T cells *in vitro*. (**A-B**) Immunoblot of Hs578T cells CRISPR-Cas9-mediated knock-out for ROSA26 (control), ezrin (EZR), radixin (RDX) and moesin (MSN) and treated with 100 nM U46619 for 5 min (**A**). p-ERM over actin signals were quantified and normalized to ROSA26 treated with vehicle (**B**). (C) Schematic representation of *in vitro* 2D migration experimental protocol. (**D-E**) Quantification of cell velocity during 2D migration of Hs578T cells following CRISPR- Cas9-mediated knock-out of ROSA26 (control) or MSN using two independent sgRNAs (**D**), and double knock-out of SLK and LOK (**E**), treated with vehicle or 10 nM U46619 for 6 hours. (**F**) Schematic representation of *in vitro* 3D invasive migration experimental protocol. (**G-H**) Quantifications of cell velocity during 3D invasive cell migration of Hs578T cells following CRISPR-Cas9-mediated knock-out of ROSA26 (control) or MSN using two independent sgRNAs (**G**), and double knock-out of SLK and LOK (**H**), treated with vehicle or 10 nM U46619 for 24 hours. *Immunoblot (**A**) is representative of three independent experiments. P-ERM quantifications (**B**) represents the mean +/- s.d. of three independent experiments*. *Velocity quantifications (**D-E**, **G-H**) represent the mean +/- s.d. of three independent experiments. Dots represent independent experiments (**B**) or individual cells (**D-E**, **G-H**). P values were calculated using Holm-Sidak’s multiple comparisons test with a single pooled variance (**D-E**, **G-H**) except for comparison made with normalizing condition (ROSA26 + vehicle, **B**) where a one-sample t-test was applied. *, P < 0.05. **, P < 0.01. ***, P < 0.001. ****, P < 0.0001. ns, not significant*.

To measure the motility of Hs578T cells, we used a modified version of the wound-healing scratch assay (Oris^TM^) (Fig 5C). Cells migrating into the free area were followed for 6 hours, a timescale during which ERMs remained activated by U46619 in Hs578T cells (Fig S3C- D). We found that upon TBXA2R activation, Hs578T cells migrated ∼1.5 faster than non stimulated cells (Fig 5D-E). The deletion of moesin by CRISPR-Cas9 using two independent single guide RNA (Fig S3E-F) significantly slowed down the spontaneous cell migration compared to controls (Fig 5D), confirming the importance of ERMs for cell migration (Arpin et al., 2011). Furthermore, we found that the TBXA2R-promoted increase of Hs578T cell migration is an ERM-dependent process, as demonstrated by the lack of increased motility in the two independent Δmoesin Hs578T cell lines following activation of this GPCR (Fig 5D). This finding underscores the importance of moesin in mediating the effect of TBXA2R on Hs578T cell migration. Confirming this, the depletion of moesin by shRNA resulted in the same cell motility defects downstream of TBXA2R activation as the one observed in the Δmoesin cells (Fig S3G). We also found that SLK and LOK double knockout, completely abrogated the TBXA2R-promoted migration confirming that they are the kinases activating ERMs downstream of TBXA2R and demonstrating their essential role in the migration resulting from activation of this GPCR (Fig 5E).

To investigate the role of the TBXA2R-ERM signaling axis during cell invasion, Hs578T cells were embedded in an extracellular collagen matrix (ECM) using the wound-healing scratch assay (Oris^TM^) (Fig 5F). Cells invading the free zone filed with ECM, using serum as a chemoattractant, were then tracked. This revealed that activation of TBXA2R by U46619 increases the speed of Hs578T cell invasion into the ECM by ∼1.5 fold (Fig 5G-H). As observed for 2D cell migration, moesin and SLK/LOK were necessary for TBXA2R to potentiate cell invasion into the ECM. Indeed, moesin or SLK/LOK knockouts prevented TBXA2R from potentiating invasion of TNBC cells (Fig 5G-H). Our findings establish that activation of ERMs via SLK/LOK downstream of TBXA2R is a key mechanism by which this GPCR promotes cell migration and invasion in TNBC cells.

Finally, we assessed whether TBXA2R signaling could potentiate TNBC metastasis through ERM activation *in vivo*. To this end, we used a tail vein metastasis assay in immunodeficient NSG mice treated or not with the TBXA2R agonist U46619 (Fig 6A). Twenty-four days after injection, the size of liver tumor nodules and the liver tumor load was compared between control Hs578T cells (ROSA26) and the two Δmoesin cell lines. Administration of U46619 was well tolerated and we did not observe any change in the body weight or weight of the liver in control mice (Fig S4A-B). In mice injected with control Hs578T cells, histopathological analysis of serial liver sections revealed that U46619 increases the size of metastatic nodules along with the liver tumor burden without affecting the number of nodules per animal (Fig 6B-D and S4C). Activation of TBXA2R also resulted in a significant increase in the percentage of mice exhibiting a tumor metastatic burden of more than 10% of liver surface, from 25% (2 out of 8 mice) to 60% (6 out of 10 mice), confirming the crucial role of this GPCR in metastatic progression (Fig 6D). Our findings also showed that mice injected with Δmoesin Hs578T cells presented fewer metastatic nodules in their livers compared to those injected with control cells (Fig S4C). In contrast to mice injected with control Hs578T cells, treatment with U46619 did not increase the average size of liver metastatic nodules nor the liver tumor load when Δmoesin Hs578T cells were injected (fig 6B-D). As a result, none of the mice injected with Δmoesin cells showed a metastatic burden of more than 10% of their liver surface (Fig 6D). Notably, the differences in nodule size or tumor load observed across the different groups were not due to an effect of U46619 on the growth of metastatic cells. Indeed, we found that U46619 did not affect the proliferation of control or Δmoesin cells *in vitro* (Fig S4D) nor *in vivo* since the Ki-67 proliferative index of these cells was similar in the liver of every animal (Fig 6E-F). Together, these observations indicate that TBXA2R promotes metastatic progression in a moesin-dependent manner.

**FIGURE 6:**
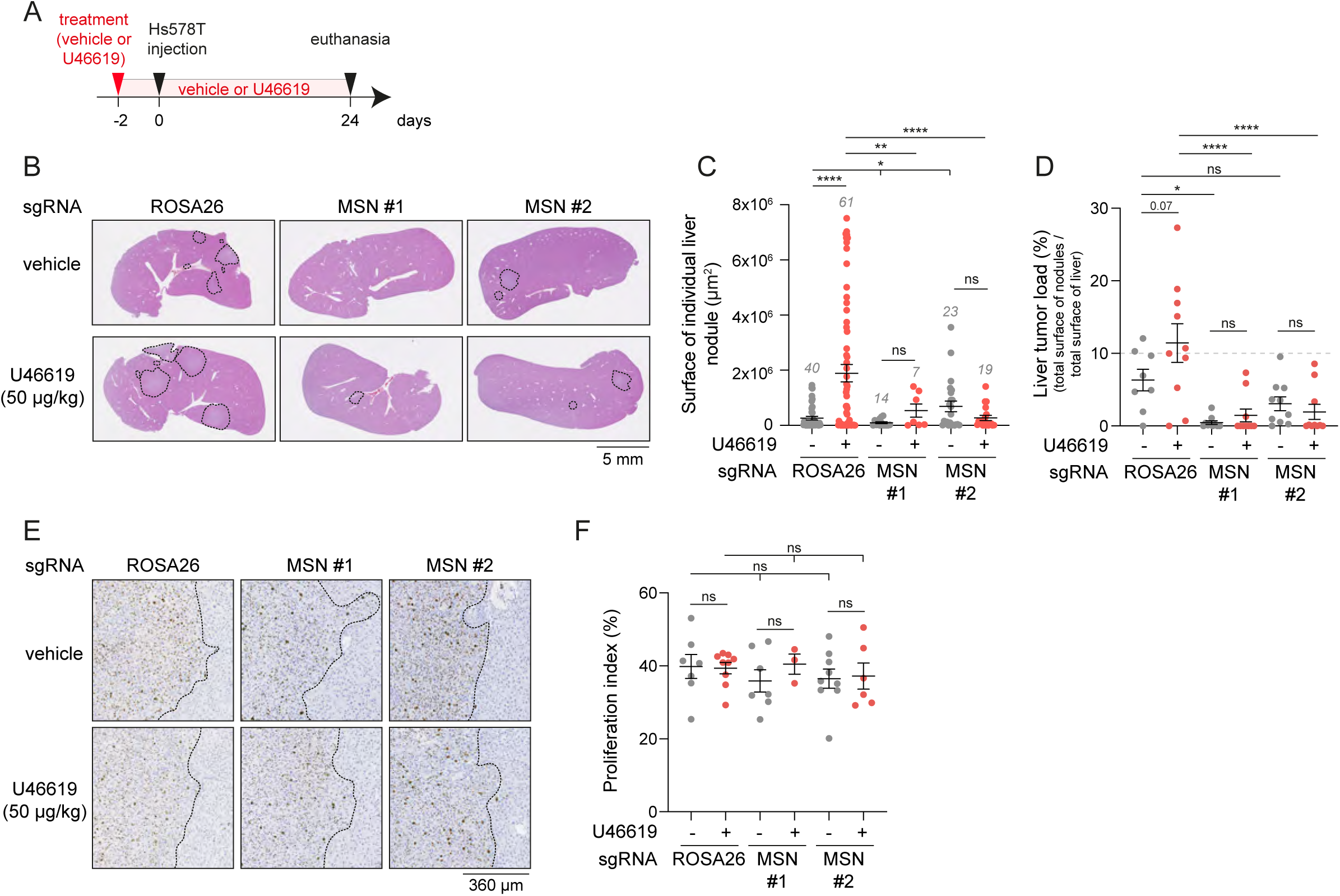
TBXA2R-induced metastasis of TNBC cells is mediated by Moesin *in vivo*. (**A**) Schematic representation of the *in vivo* tail vein forced metastasis assay. NSG mice were injected with Hs578T cells CRISPR-Cas9-mediated knock-out for ROSA26 (control) or MSN using two independent sgRNAs. From two days before injection until euthanasia, mice were treated with vehicle or U46619 administered at 50 µg/kg through drinking water. n = 8 mice for ROSA26 + vehicle, n = 10 mice for all other condition. (**B-D**) Characterization of liver tumor nodules in mice treated as detailed in panel A. (**B**) Representative H&E staining of liver sections. Black lines highlight tumor nodules. (**C-D**) The surface of individual tumor nodules was measured using NDP view 2 (Hamamatsu Photonics) on H&E-stained liver sections and represented as individual tumor nodule (**C**) or represented as a ratio of the total surface of nodules over total surface of liver per animal (expressed in %) (**D**). Quantifications represent the mean +/- s.e.m of 8 or 10 animals. (**E-F**) Representative IHC staining for Ki-67 in liver sections from mice described in **A**. Black lines demarcates the tumor (left) from the adjacent normal tissue (right) (**E**). The proliferation index (Ki-67^+^ cells expressed in %) was then quantified (**F**). Quantifications represent the mean +/- s.e.m of 8 or 10 animals. *Dots represent individual tumor nodules (**C**) or individual animals (**D, F**). P values were calculated using Welch’s t-test because of unequal sample sizes (**C**) or Holm-Sidak’s multiple comparisons test with a single pooled variance (**D, F**). *, P < 0.05. **, P < 0.01*. *\*\*\*\*, P < 0.0001. ns, not significant*.

## DISCUSSION

Our study sheds light on the involvement of a GPCR-ERM signaling pathway in cancer cell biology. We have discovered that the TBXA2R activates proteins of the ERM family, thereby enhancing cancer cell motility and invasion *in vitro,* as well as metastatic colonization in a mouse xenograft model. This signaling pathway is mediated through the Gα_q/11_ and Gα_12/13_ subfamilies, which both converge on RhoA. In turn, RhoA activates kinases of the SLK family that promote ERM activation.

Previous studies have identified ROCK, SLK, and LOK kinases as direct effectors of RhoA (Bagci et al., 2020; Matsui et al., 1996) and activators of ERMs (Belkina et al., 2009; Machicoane et al., 2014; Matsui et al., 1998; Viswanatha et al., 2012). In our experiments with HEK293T and Hs578T cells, we found that SLK and LOK, but not ROCK, are responsible for ERM activation, both under steady-state conditions and upon TBXA2R activation.

Recently, we reported that RhoA activates SLK to drive metaphase cell rounding by activating ERMs at the mitotic cortex (Leguay et al., 2022). We found that two different Rho-GEF, GEF-H1 and Ect2 activate RhoA at mitotic entry. Here we showed that ERM activation depends on RhoA downstream of Gα_q/11_ and Gα_12/13_ subfamilies. While Gα_q/11_ inhibition alone did not prevent ERM activation downstream of TBXA2R in HEK293T and TNBC cells, genetic depletion of Gα_12/13_ partially prevented this activation. It is only when both Gα subfamilies are inhibited that TBXA2R signaling fails to activate ERMs. This indicates that whereas Gα_12/13_ is the main Gα subfamily engaged by TBXA2R to activate ERMs, Gα_q/11_ can partially compensate for the loss of Gα_12/13_. Several Rho-GEF such as p63RhoGEF, p115RhoGEF, pDZ-RhoGEF, GEF-H1, and LARG (Lutz et al., 2007; Meiri et al., 2014; Nakahata, 2008; Sah et al., 2000; van Unen et al., 2016; Wikstrom et al., 2008) were shown to be activated by Gα_12/13_ or Gα_q/11_ and represent possible links between these G proteins and RhoA activation.

Although GPCRs and ERMs have been extensively studied, only a few studies have reported a link between these two protein families. For instance, activation of the muscarinic M1 receptor (M1R) or the β2-adrenergic receptor (β2AR) by their respective agonists was shown to promote ezrin activation through a mechanism involving the G protein-coupled receptor kinase 2 (Cant and Pitcher, 2005). Thrombin that activates PAR1, PAR3, and PAR4 was also reported to induce the recruitment of radixin at the plasma membrane of endothelial cells upon direct binding of this ERM to Gα_13_ (Vaiskunaite et al., 2000). Lysophosphatidic acid (LPA), which activates LPARs_1-6_ also promotes the activation of moesin in platelets (Nakamura et al., 1995; Retzer and Essler, 2000; Shcherbina et al., 1999). More recently, LPA activation of LPAR_1_ and LPAR_2_ was found to stimulate the phosphorylation of ERMs in the ovarian cancer cell line OVCAR-3 (Park et al., 2018). Interestingly, expression of ezrin^T567A^, a non-phosphorylatable mutant of the regulatory threonine that acts as a dominant negative form of ERMs, reduced the migration of OVCAR-3 cells toward LPA, used as a chemoattractant.

Our study establishes the signaling cascade that connects a GPCR, TBXA2R, to ERM activation, which ultimately promotes TNBC cell migration, invasion and metastatic colonization. While it is currently unclear whether alternative mechanisms proposed for other GPCRs, such as the involvement of GRK2 (Cant and Pitcher, 2005) or direct binding of ERM to Gα_13_ (Vaiskunaite et al., 2000) contribute to ERM activation downstream of TBXA2R, our findings demonstrated that kinases of the SLK family are the only kinases needed for ERM activation by TBXA2R. Therefore, any potential contribution from other mechanisms would need to occur upstream of these two kinases. It is also unclear whether Gα_q/11_ and/or Gα_12/13_ are the only G proteins that mediate the activation of ERM downstream of the other GPCRs found to activate ERMs. However, it is noteworthy that M1R, β2AR, Thrombin-activated PARs, as well as LPAR_1/2_, were found to activate either Gα_q/11_ or Gα_12/13_ or both (Avet et al., 2022).

The molecular basis underlying metastasis is still incompletely understood, and the mechanisms that prompt cancer cells to disseminate to secondary organs are very diverse. These mechanisms depend on genetic alterations of cancer cells, but they also rely on the interaction of cancer cells with the stroma (Welch and Hurst, 2019). Thromboxane A2 (TXA2), the natural ligand of TBXA2R, was found to promote metastasis in colorectal (Guillem-Llobat et al., 2016), lung (Lucotti et al., 2019), prostate (Nie et al., 2004) and breast (Ekambaram et al., 2011) cancers, opening the possibility that TBXA2R can be targeted to prevent metastasis. Supporting this hypothesis, Ifetroban (CPI211), a potent and selective antagonist of TBXA2R, was recently shown to reduce the metastatic burden of several cancer cell lines, including TNBC cells, in xenograft mouse models (Werfel et al., 2020). Ifetroban is currently being evaluated in a clinical trial (NCT03694249) as a potential treatment for high-risk solid tumors that are prone to metastatic recurrence. The rationale of this trial is that Ifetroban inhibits TBXA2R on platelets, which reduces their aggregation on cancer cells, ultimately preventing platelet- assisted metastasis (Lucotti et al., 2019; Werfel et al., 2020).

Our study identifies a distinct mechanism by which TBXA2R could be targeted to block metastasis; activation of TBXA2R on TNBC cells can increase their motility, invasion, and metastatic potential by activating ERMs. Overexpression of essential components of this pathway could upregulate this TBXA2R-ERM signaling axis. Supporting this notion, we found that TBXA2R expression correlates with phosphorylation and activation of ERMs in TNBC patient samples. Since TBXA2R signaling is a potent activator of ERMs, activation of this GPCR by its natural ligand, TXA2, could also promote metastasis by activating ERMs. Identifying therapeutic targets for the treatment of metastatic disease is challenging due to its complexity, and so far, no effective anti-metastatic drugs have been approved for clinical use (Anderson et al., 2019). However, the discovery that TBXA2R activates ERMs through Gα_12/13_ and Gα_q/11_ suggests that other GPCRs that engage one of these two Gα subfamilies could also promote metastasis by activating ERMs. Given that several GPCR antagonists are already approved drugs or have advanced to clinical trials for other indications, repurposing them for anti-metastatic treatment could represent an appealing avenue that could accelerate the discovery of new anti-metastatic treatments. Moreover, our characterization of the TBXA2R-ERM signaling pathway reveals novel potential therapeutic targets to prevent metastasis. While ERMs are already promising therapeutic targets (Clucas and Valderrama, 2014; Ghaffari et al., 2019; Hoskin et al., 2019; Ren and Khanna, 2014), their activating kinases, SLK and LOK, which have not been explicitly considered as targets yet, could also represent promising targets to prevent metastatic dissemination and outgrowth.

## ACKNOWLEDGMENT

This work has been supported by a project Grant from the CIHR (175193) to S.C. and M.B., a Foundation Grant from the CIHR (148431) to M.B. and a Cancer Research Society grant (25388) to S.M. K.L held a doctoral scholarship from Institute for Research in Immunology and Cancer (IRIC) and from Montreal University’s Molecular Biology Program as well as a Études Supérieures et Postdoctorales (ESP) studentship from Montreal University. O.N. holds a master and doctoral scholarship from the Mission Universitaire de Tunisie en Amérique du Nord, from Montreal University’s Molecular Biology Program as well as from IRIC. C.T. held a doctoral studentship from the Fonds de recherche Santé Québec. M.B. held the Canada Research Chair in Signal Transduction and Molecular Pharmacology. The authors are grateful to Dr. Asuka Inoue for providing the ΔGα_q/11_ and ΔGα_12/13_ HEK293 cells. The authors declare no competing financial interests.

## AUTHOR CONTRIBUTIONS

Management of the project: K.L., C.L.G, M.B, SC. Conceptualization: K.L., C.L.G, S.M, M.B, SC. Investigation: K.L., O.N, Y.Y.H, M.H, C.T, E.M.G, H.K. Formal Analysis: K.L., O.N, H.K, C.L.G, S.M, M.B, S.C. Resources: L.G. Writing – Original Draft: K.L, M.B, S.C. Writing – Review and Editing: K.L, C.L.G, S.M, M.B, S.C. Visualization: K.L, H.K, S.C. Funding acquisition: S.M, M.B, S.C.

## MATERIAL AND METHODS

### Reagents and inhibitors

Coelenterazine 400a (Deep Blue C) was purchased from NanoLight Technology (#340). U46619 and SQ-29548 were purchased from Cayman Chemical (#16450 and #19025, respectively). Calyculin A and Y-27632 ROCK inhibitors were purchased from Sigma (#C5552 and #688000, respectively). Staurosporine was purchased from ApexBio (#A8192). Rho inhibitor I (C3 transferase) was purchased from Cytoskeleton (#CT04). YM- 254890 was purchased from FUJIFILM Wako Pure Chemical Corporation (#257-00631).

### DNA constructs

The cDNA for TBXA2R was previously described (Parent et al., 1999). (MyrPB-)ezrin-, radixin-, moesin-rLucII, and rGFP-CAAX constructs were previously described (Leguay, JCS 2021). MISSION shRNA constructs were obtained in pLKO.1-puro vectors from Sigma: EZR (TRCN0000062459), RDX (TRCN0000062434), MSN (TRCN0000062412). FlexiTube siRNA were obtained from Qiagen: SLK#1 (SI00107723), SLK#2 (SI04438350), LOK (SI02224054).

### Cell culture and transfection

HEK293T human kidney cells, Hs578T and MDA-MB-231 TNBC cells were cultured in Dulbecco’s modified Eagle’s medium (DMEM; 4.5 g/L D-Glucose, L-Glutamine, 110 mg/L sodium pyruvate; Thermofisher #11995073), MB-468 and BT-549 TNBC cells in RPMI 1640 medium (Invitrogen) at 37°C with 5% CO_2_. DMEM and RPMI were supplemented with 10% fetal bovine serum (FBS; Life Invitrogen #12483020) and 1% penicillin-streptomycin antibiotics (ThermoFisher #15070063). MDA-MB-157 cells were cultured in Leibovitz’s L- 15 medium (Cedarlane), supplemented with 15% FBS and 1% penicillin-streptomycin, and maintained at 37°C without CO_2_.

HEK293T cells knock-out for G_q/11_ or G_12/13_ were obtained from Asuka Inoue (Tohoku University, Japan). Hs578T cell clones knock-out for ezrin, radixin, moesin, or SLK/LOK were obtained by CRISPR-Cas9, followed by clonal dilution after selection with 2 µg/ml puromycin (EMD Millipore #540222) for 48h. The following sequence guides were inserted in plentiCRISPR-v2 (addgene #52961) digested with BsmBI (New England Biolabs #R0739): EZR (CTGAGCGGCTGATCCCTCAA), RDX (GTTTCACTTACCGCTGGGGT), MSN#1 (GAGACAAGTTGCTCCCGCAG), MSN#2 (AAGCTTACCTGAGCATGCCA), SLK (CAATTTGATATCTCCATCTA) and LOK (CATGATTGAGTTCTGTCCAG).

For ebBRET experiments, HEK293T cells were transfected using linear polyethyleneimine (PEI, Alfa Aesar #43896) as previously described (Leguay et al., 2021). Loss of function experiments were performed by transfecting siRNA using lipofectamine RNAiMAX (Thermofisher #13778075) as prescribed by the manufacturer or by infecting cells with lentiviral particles in DMEM supplemented with 10% FBS and 5 µg/ml polybrene (Sigma #H9268) for 48h followed by a 48h-treatment with 2 µg/ml puromycin.

### ebBRET measurement

ebBRET measurements were performed as previously described (Leguay et al., 2021). Briefly, 48h after transfection, HEK293T cells were washed with Hank’s balanced salt solution (HBSS, ThermoFisher #14065056) and incubated for 5 min with 2.5 µM coelenterazine 400a diluted in HBSS. ebBRET signals were monitored using a Tecan Infinite 200 PRO multifunctional microplate reader (Tecan) equipped with BLUE1 (370- 480 nm; donor) and GREEN1 (520-570 nm; acceptor) filters. ebBRET signals were calculated as a ratio by dividing the acceptor emission value by the donor emission value.

### Immunoblotting

Following the indicated treatment, cells were washed with ice-cold phosphate-buffered saline (PBS) and lysed in TLB buffer (40 mM HEPES, 1 mM EDTA, 120 mM NaCl, 10 mM NaPPi, 10% glycerol, 1% Triton X-100, 0.1% SDS) supplemented with both phosphatase and protease inhibitors (phosphatase inhibitor cocktail (PIC, Sigma #P2850), 1 mM sodium orthovanadate (Na_3_VO_4_, Sigma #S6508), 5 mM β-glycerophosphate (Sigma #G6251), 1 mM phenylmethylsulfonyl fluoride (PMSF, Sigma #P7626) and anti-protease cocktail (Sigma #4693132001)). Samples were then denatured in sample buffer (200 mM Tris-HCl 1M pH6.8, 8% SDS, 0.4% bromophenol blue, 40% glycerol, and 412 mM β-mercaptoethanol) and resolved by SDS-PAGE followed by transfer to nitrocellulose membranes (pore 0.2μm, VWR #27376-991). Membranes were blocked in TBS-Tween (25 mM Tris-HCl pH 8, 125 mM NaCl, 0.1% Tween 20) supplemented with 2% BSA for one hour before overnight incubation with primary antibodies at 4°C. Primary antibodies were: rabbit anti-p-ERM (1:5000, Roubinet 2011), rabbit-anti ERM (1:1000, Cell Signaling #3142), rabbit anti-Ezrin (1:1000, Cell Signaling #3145), rabbit anti-Radixin (1:1000, Cell Signaling #2636), rabbit anti-Moesin (1:1000, Cell Signaling #3150), mouse anti-Actin (1:5000, Sigma #MAB1501), rabbit anti-SLK (1:1000, Cerdalane #A300-499A) and rabbit anti-LOK (1:5000, Abcam #ab70484). Washed membranes were then incubated for 1h at room temperature with secondary antibodies: goat anti-rabbit-IgG HRP (1:10000, Santa Cruz biotechnology #sc-2004) and goat anti-mouse-IgG HRP (1:10000, Santa Cruz Biotechnology #sc-516102). Protein detection was finally performed using Amersham ECL Western Blotting detection reagent (GE Healthcare #CA95038-564L).

### In vitro kinase assay

In vitro kinase assays were performed as previously described (Leguay et al., 2022). Briefly, HEK293T cells treated with U46619 or its vehicle were lysed in TLB buffer supplemented with phosphatase and protease inhibitors. Endogenous SLK was then immunoprecipitated from cell lysate using rabbit anti-SLK antibody (Bethyl #A300-499A) incubated for 1h at 4°C, followed by incubation with protein A-sepharose beads (GE Healthcare #GE17-0780-01) for 2h at 4°C. Beads were then extensively washed and resuspended in kinase reaction buffer (KRB; 50 mM Tris-HCl pH 7.5, 100 mM NaCl, 6 mM MgCl_2,_ and 1 mM MnCl_2_) supplemented with both phosphatase and protease inhibitors, 2 mM DTT, 50 µM ATP and purified recombinant GST-Ezrin^479-585^ from BL21 bacteria. Kinase assay was finally performed for 30 min at 30°C, and proteins were denatured with sample buffer.

### Immunofluorescence

The day before the experiment, cells were plated on glass coverslips (Marienfeld #0115200) and incubated at 37°C with 5% CO_2_ overnight. Cells were then washed with PBS and fixed with 10% trichloroacetic acid (Sigma #T0699) for 10 min at room temperature, followed by extensive washes with TBS (20mM Tris-HCl pH 7.5, 154 mM NaCl, 2mM EGTA, 2mM MgCl_2_). Cells were then permeabilized with 0.02% saponin (Amresco #0163) diluted in TBS supplemented with 2% BSA (Bioshop #ALB001.250) for 1h. Cells were then incubated overnight with rabbit anti-p-ERM primary antibody (1:5000, Roubinet, 2011) followed by goat anti-rabbit Alexa Fluor 488-conjugated secondary antibody (1:200, Invitrogen #A11070) for 1h. Glass coverslips were finally mounted in Vectashield medium with DAPI (Vector Laboratories #H-1200), and images were acquired using an LM700 confocal microscope (Zeiss) with a 63x objective.

### Cell migration and invasion assay

Migration experiments were performed using the Oris^TM^ Cell Migration assay (Platypus technologies) as the manufacturer prescribed. Briefly, Hs578T cells were plated in DMEM supplemented with 10% FBS in a 96-well plate with stoppers creating a central cell-free detection zone. The next day, stoppers were removed, and media was replaced with DMEM supplemented with 10% FBS and 10 nM U46619 or vehicle. Cells were then allowed to migrate into the free area.

Invasion experiments were performed using the Oris^TM^ Cell Invasion assay (Platypus technologies) as prescribed by the manufacturer. Briefly, Hs578T were plated embedded in DMEM supplemented with 2 mg/ml collagen type I (rat tail) (Fisher scientific #CB354249) in a 96-well plate with stoppers creating a central cell-free detection zone. After collagen polymerization, stoppers were removed, and DMEM supplemented with 10% FBS and 2 mg/ml collagen type I was added to the central cell-free detection zone. After collagen polymerization, DMEM supplemented with 10 nM U46619, or vehicle was added on top of collagen. Cells could then invade into the free area.

Migration and invasion of cells into the free zone were recorded by contrast video microscopy using LSM700 confocal microscope (Zeiss) with 20x objective for 6 or 24 hours, respectively. Individual cells were manually tracked using the Manual Tracking plugin on ImageJ software (NIH). Cell velocity was determined as a ratio by dividing the total distance v over time. Representations of cell migration were obtained using Chemotaxis and Migration Tool 2.0 (Ibidi).

### Immunohistochemistry

Tissue microarray was obtained as previously described (Yousef et al., 2014). Samples were obtained from the *Centre Hospitalier de l’Université de Montréal* (CHUM) after approval by the research ethical committee (Comité d’éthique de la recherche du CHUM CENTRE DE RECHERCHE, Approval No. SL 05.019).

Livers excised from mice were fixed in 10% formalin, embedded in paraffin, and sliced into 4-µm sections. Tissue sections were mounted on glass slides and stained with hematoxylin and eosin (H&E) using conventional protocols. Immunohistochemical assays were performed on the Bond RX Stainer (Leica Biosystems, Buffalo Grove, IL, USA) according to the manufacturer’s instructions. Briefly, slides were deparaffinized, and antigen recovery was conducted by heat-induced epitope retrieval with standard CC1 and CC2 solutions (Ventana Medical Systems, Tucson, AZ). Sections were then incubated with primary antibody for 30 min. Primary antibodies used are the following: p-ERM (1:600, (Roubinet et al., 2011)), TBXA2R (1:100, Cayman chemicals #10004452) and Ki-67 (1:150, Biocare Medical #CRM325B). Bound primary antibodies were detected using peroxidase polymer (HRP-DAB) for 15 min. The stained slides were next subjected to digital slide scanning that converts glass slides into high-resolution digital data by high-speed scanning using the NanoZoomer Digital Pathology (NDP) 2.0-HT digital slide scanner (Hamamatsu, Japan). Signal intensities for p-ERM and TBXA2R were quantified at the plasma membrane using Image J software (NIH). Quantification of Ki-67-positive cells was performed using Visiomorph software (Visiopharm).

### In vivo metastasis assay

Hs578T cells (5.0 x 10^4^) were resuspended in 100 µl PBS and injected into the tail vein of NSG mice. Two days before cell injection and during the whole experiment, NSG mice were treated with U46619 (50 µg/kg) or vehicle (DMSO) administered through drinking water containing 0.1% aspartame (TCI chemicals #A0997), given ad libitum and changed every second day. The mice were then euthanized 24 days after cell injection in a CO_2_ chamber. At the time of euthanasia, the liver was photographed and weighed, and the number of tumors was manually assessed under a stereo microscope. The left lobe was then processed for histological analysis.

### Cell proliferation assay

Cell proliferation was measured using MTT. Briefly, Hs578T cell lines were seeded into 96- well plates in quadruplicate and cultured for the durations indicated in the figure legend. The medium was then replaced with fresh medium supplemented with 1.1 mM MTT (Cayman chemical #21795), and cells were incubated at 37°C for 4h. Dissolution of formazan was then performed by incubating cells with 67% DMSO at 37°C for 10 min. Cell proliferation was assessed by reading the absorbance at 540 nm using a Tecan Infinite 200 PRO multifunctional microplate reader (Tecan).

### Data analysis

All quantifications were performed using ImageJ software (NIH) and analyzed using GraphPad PRISM software (GraphPad Software, La Jolla, CA, USA). Microscopy images were prepared using Image J software (NIH) and Photoshop (Adobe).

For cancer survival analysis, we used gene expression data from triple-negative breast cancer (TNBC) and normal breast tissues obtained from The Cancer Genome Atlas (TCGA). The TCGA-BRCA RNA-seq dataset was downloaded from the Broad GDAC Firehose (https://gdac.broadinstitute.org/). For survival analyses, we calculated the mean expression of *TBXA2R* and *MSN* and categorized the samples into groups based on the upper and lower tertiles. Overall survival was analyzed using the Kaplan–Meier method, and differences in survival rates were compared using the log-rank test in GraphPad PRISM software.

---

**SUPPLEMENTARY FIGURE 1.**
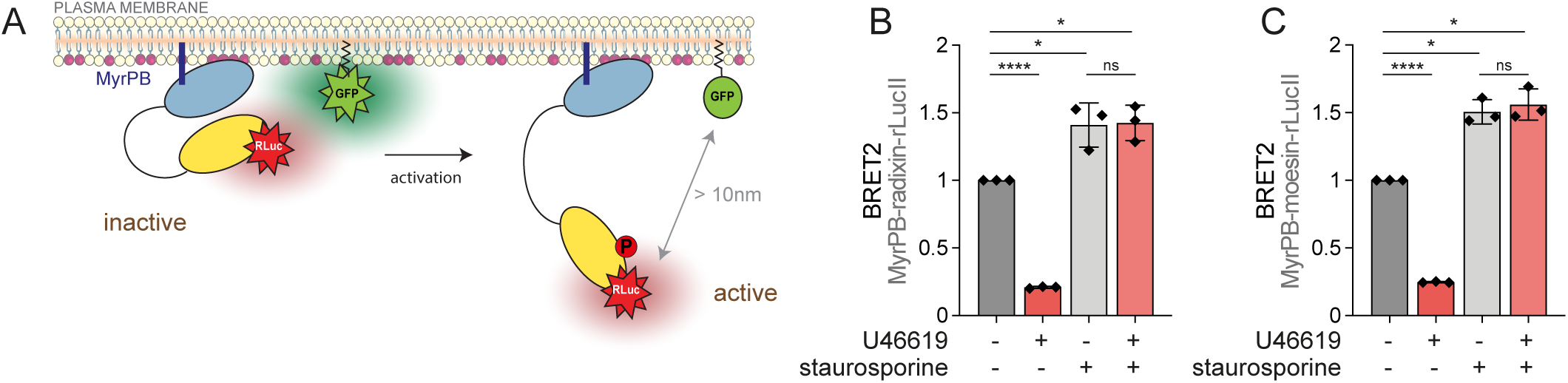
(**A**) Schematic representation of MyrPB-E,R,M-rLucII ebBRET biosensors (adapted from (Leguay et al., 2021)). (**B-C**) ebBRET signals measured in HEK293T-TBXA2R cells expressing MyrPB-radixin-rLucII (**B**) or MyrPB-moesin-rLucII (**C**) biosensor and treated with 100 nM U46619 for 5 min and/or 100 nM staurosporine for 30 min. *ebBRET signals (**B**-**C**) represent the mean +/- s.d. of three independent experiments. Dots represent independent experiments (**B**-**C**). P values were calculated using a two-tailed paired t-test (**B**-**C**) except for comparison made with normalizing condition (vehicle) where a one-sample t-test was applied. *, P < 0.05. ****, P < 0.0001. ns, not significant*.

**SUPPLEMENTARY FIGURE 2.**
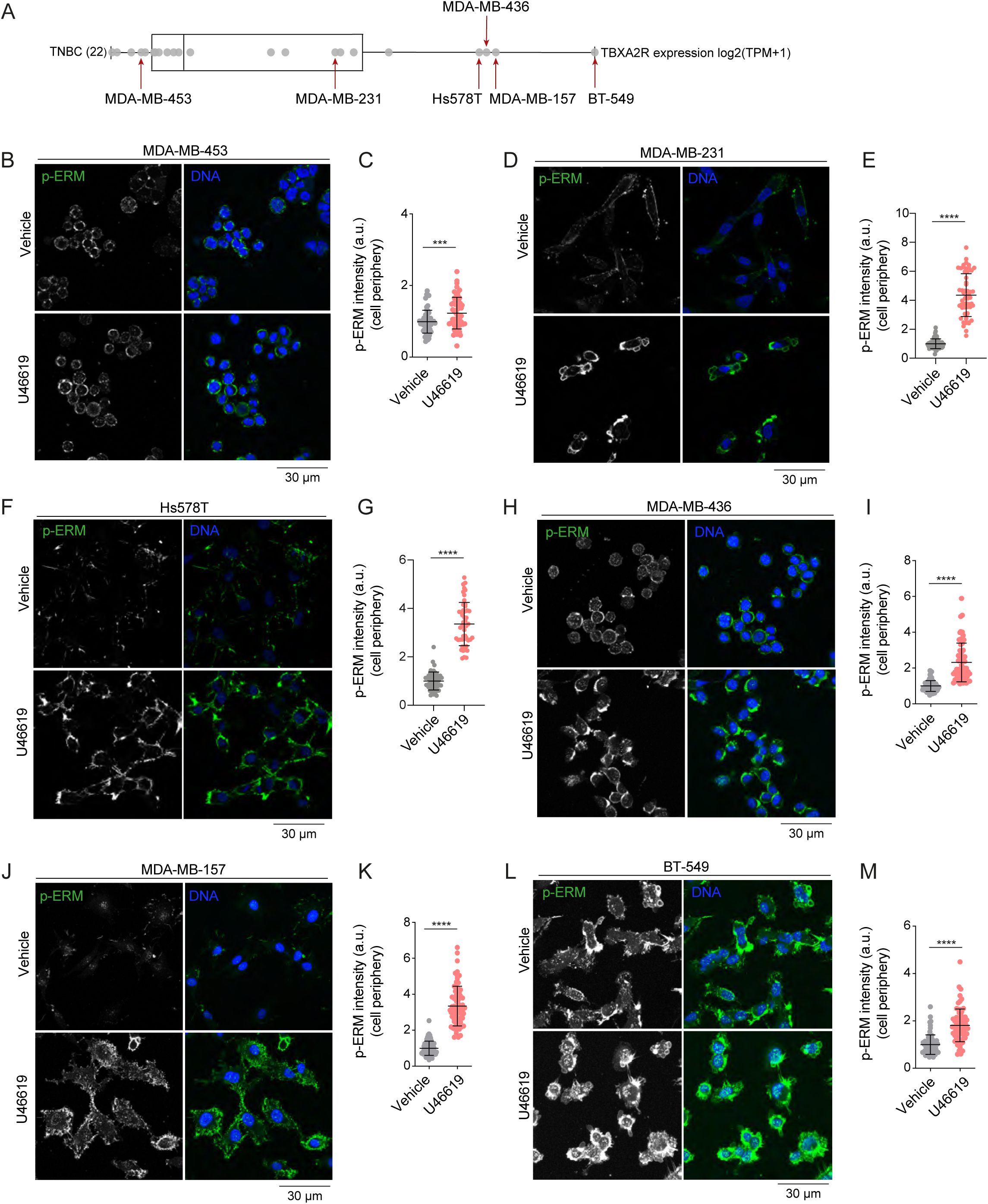
(**A)** Dot plot showing TBXA2R mRNA expression levels across various TNBC cell lines from the DepMap dataset. Each dot represents a specific TNBC cell line. The x-axis lists the cell lines ordered by increasing TBXA2R expression, measured as log2(TPM+1). Cell lines on the left exhibit lower expression levels, while those on the right exhibit higher expression levels. (**B-L**) Immunofluorescence of MDA-MB-453 (**B**), MDA-MB-231 (**D**), Hs578T (**F**), MDA-MB -436 (**H**), MDA-MB-157 (**J**), and BT-549 (**L**) cells treated with vehicle or 100 nM U46619 for 5 minutes. Peripheral p-ERM staining was quantified and normalized to vehicle-treated cells: (**C**) MDA-MB-453, (**E**) MDA-MB-231, (**G**) Hs578T, (**I**) MDA-MB-436, (**K**) MDA-MB-157, (**M**) BT-549. *Immunofluorescences are representative of three independent experiments. P-ERM quantifications represent the mean +/- s.d. of three independent experiments. Dots represent individual cells (**D**). P values were calculated using a one-sample t-test applied*. *\*\*\*\*, P < 0.0001. ***, P < 0.001. ns, not significant*.

**SUPPLEMENTARY FIGURE 3.**
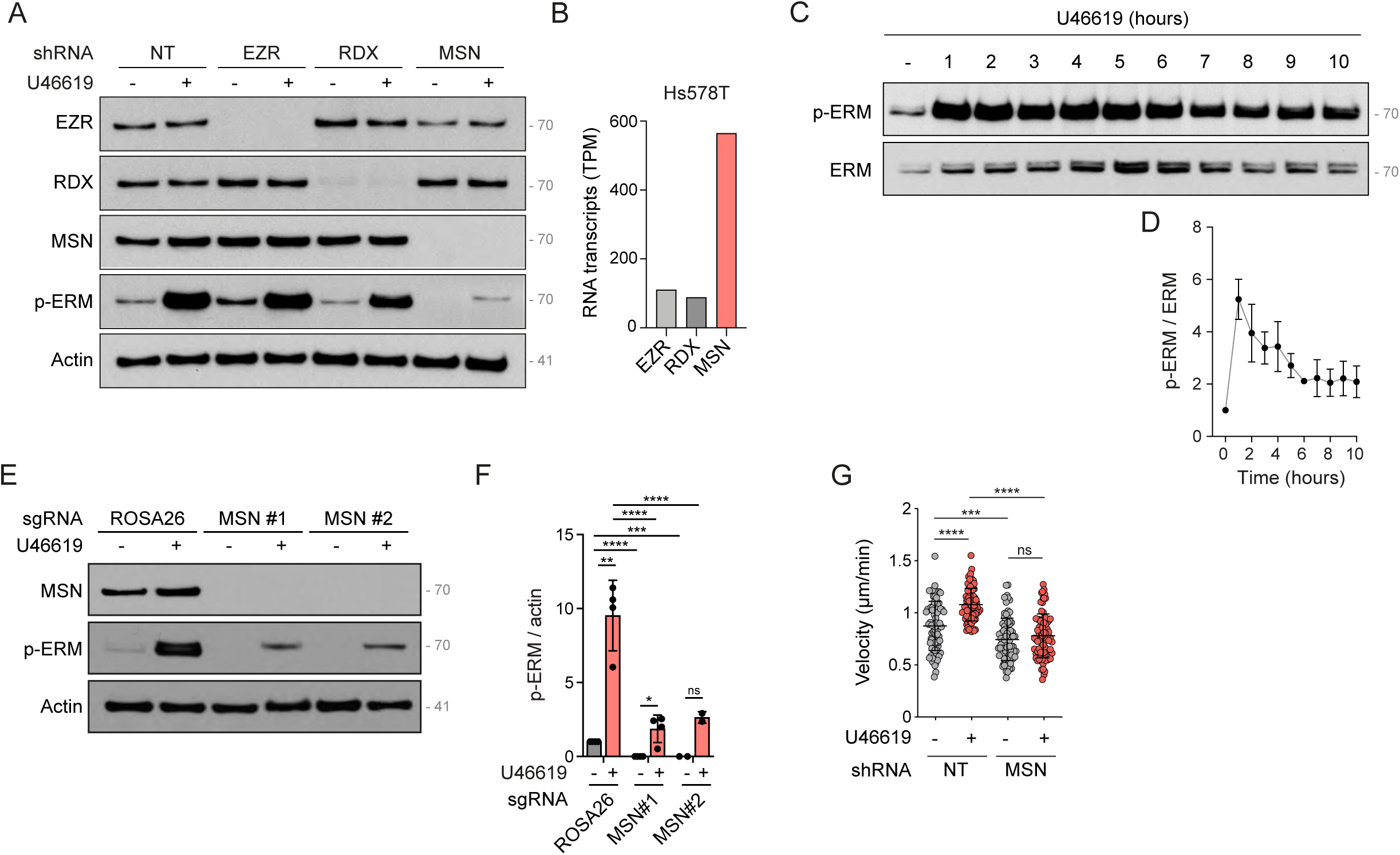
(**A**) Immunoblot of Hs578T stably expressing non-target shRNA (NT) or shRNA targeting EZR, RDX, or MSN. (**B**) mRNA expression of EZR, RDX, and MSN in the Hs578T cell line. Data extracted from single-cell RNAseq (Papatheodorou et al., 2020). (**C-D**) Immunoblot of Hs578T cells treated with 100 nM U46619 for the indicated time in hours (**C**). p-ERM over ERM signals were quantified and normalized to the vehicle (**D**). (**E-F**) Immunoblot of Hs578T cells knock-out for ROSA26 (control) or MSN using two independent sgRNAs and treated with 100 nM U46619 for 5 min (**E**). p-ERM over actin signals were quantified and normalized to ROSA26 + vehicle (**F**). (**G**) Quantifications of cell velocity during 2D cell migration of Hs578T cells stably expressing non-target shRNA (NT) or an shRNA targeting MSN and treated with vehicle or 10 nM U46619 for 6 hours. *Immunoblots (**A**, **C, E**) are representative of three independent experiments. Velocity quantifications (**G**) represent the mean +/- s.d. of three independent experiments. Dots represent the mean of independent experiments (**F**) or individual cells (**G**). P values were calculated using Holm-Sidak’s multiple comparisons test with a single pooled variance **, P < 0.01. ***, P < 0.001. ****, P < 0.0001. ns, not significant*.

**SUPPLEMENTARY FIGURE 4.**
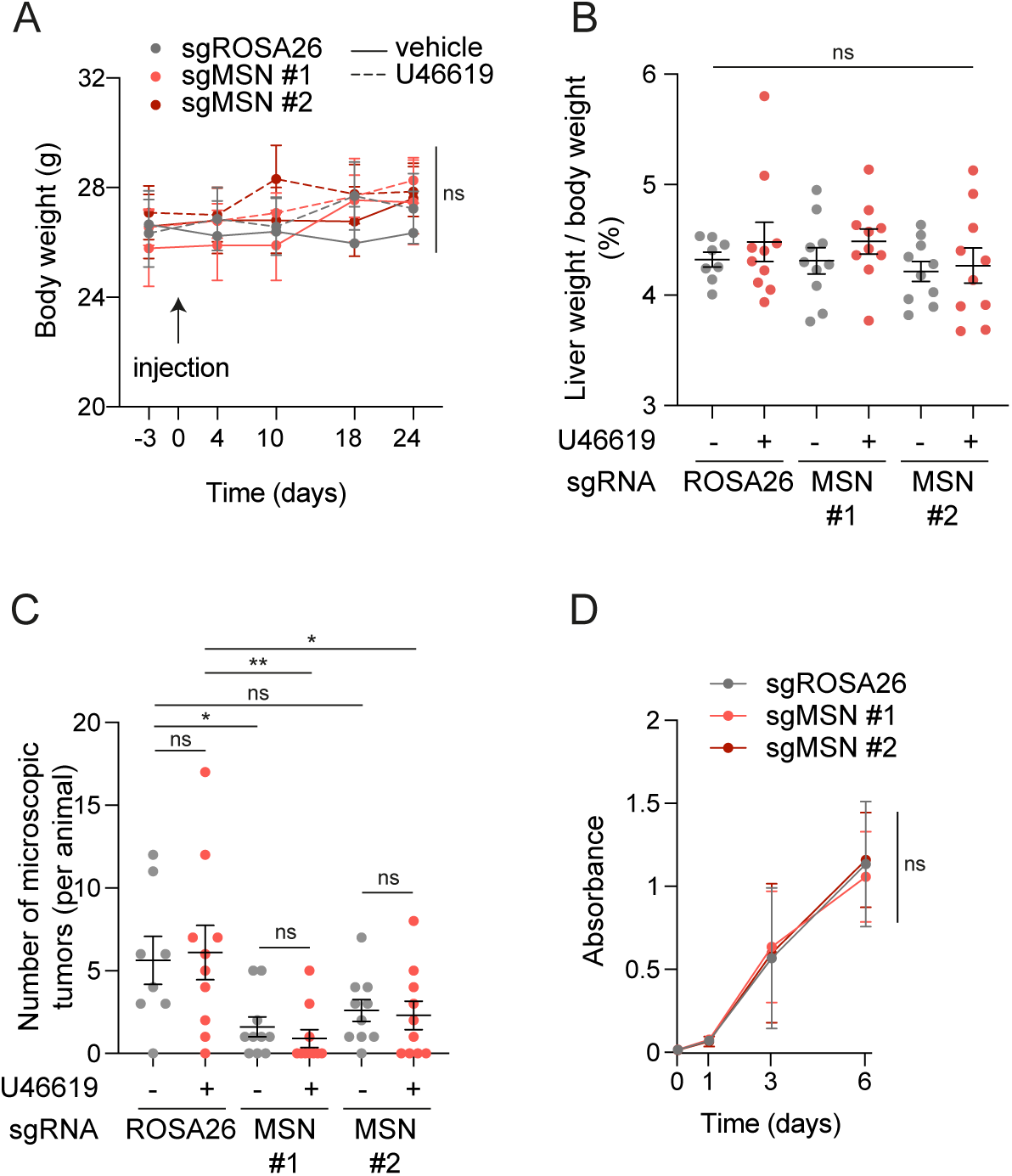
(**A**) Measurement of body weights throughout the *in vivo* experiment. Data represent mean +/- s.d. of all animals. (**B**) Measurement of liver/body weight ratios. At the time of euthanasia, mice were weighed, and the livers were excised and weighed. Data represent mean +/- s.e.m. of all animals. (**C**) Quantification of tumor nodules observed in livers from mice described in Fig 6. (**D**) Cell proliferation of Hs578T cells knock-out for ROSA26 (control) or MSN using two independent sgRNAs measured *in vitro* for 6 days. Data represent the mean +/- s.d. of three independent experiments. *Dots represent the mean of independent animals (**A**), individual animals (**B, C**) or the mean of three independent experiments (**D**). P values were calculated using a two-tailed paired t-test (**A, D**) or using Holm-Sidak’s multiple comparisons test with a single pooled variance (**B, C**). *, P < 0.05. **, P < 0.01. ns, not significant*.

## REFERENCE

Algrain, M., Turunen, O., Vaheri, A., Louvard, D., and Arpin, M. (1993). Ezrin contains cytoskeleton and membrane binding domains accounting for its proposed role as a membrane-cytoskeletal linker. J Cell Biol 120, 129–139.

Anderson, R.L., Balasas, T., Callaghan, J., Coombes, R.C., Evans, J., Hall, J.A., Kinrade, S., Jones, D., Jones, P.S., Jones, R., et al. (2019). A framework for the development of effective anti-metastatic agents. Nat Rev Clin Oncol 16, 185–204.

Arpin, M., Chirivino, D., Naba, A., and Zwaenepoel, I. (2011). Emerging role for ERM proteins in cell adhesion and migration. Cell Adh Migr 5, 199–206.

Atwood, B.K., Lopez, J., Wager-Miller, J., Mackie, K., and Straiker, A. (2011). Expression of G protein-coupled receptors and related proteins in HEK293, AtT20, BV2, and N18 cell lines as revealed by microarray analysis. BMC genomics 12, 14.

Avet, C., Mancini, A., Breton, B., Le Gouill, C., Hauser, A.S., Normand, C., Kobayashi, H., Gross, F., Hogue, M., Lukasheva, V., et al. (2020). Selectivity Landscape of 100 Therapeutically Relevant GPCR Profiled by an Effector Translocation-Based BRET Platform. 2020.2004.2020.052027.

Avet, C., Mancini, A., Breton, B., Le Gouill, C., Hauser, A.S., Normand, C., Kobayashi, H., Gross, F., Hogue, M., Lukasheva, V., et al. (2022). Effector membrane translocation biosensors reveal G protein and betaarrestin coupling profiles of 100 therapeutically relevant GPCRs. Elife 11.

Bagci, H., Sriskandarajah, N., Robert, A., Boulais, J., Elkholi, I.E., Tran, V., Lin, Z.Y., Thibault, M.P., Dube, N., Faubert, D., et al. (2020). Mapping the proximity interaction network of the Rho-family GTPases reveals signalling pathways and regulatory mechanisms. Nat Cell Biol 22, 120–134.

Barik, G.K., Sahay, O., Paul, D., and Santra, M.K. (2022). Ezrin gone rogue in cancer progression and metastasis: An enticing therapeutic target. Biochim Biophys Acta Rev Cancer 1877, 188753.

Bartholow, T.L., Becich, M.J., Chandran, U.R., and Parwani, A.V. (2011). Immunohistochemical analysis of ezrin-radixin-moesin-binding phosphoprotein 50 in prostatic adenocarcinoma. Bmc Urol 11.

Bartova, M., Hlavaty, J., Tan, Y., Singer, C., Pohlodek, K., Luha, J., and Walter, I. (2017). Expression of ezrin and moesin in primary breast carcinoma and matched lymph node metastases. Clinical & experimental metastasis 34, 333–344.

Belkina, N.V., Liu, Y., Hao, J.J., Karasuyama, H., and Shaw, S. (2009). LOK is a major ERM kinase in resting lymphocytes and regulates cytoskeletal rearrangement through ERM phosphorylation. Proc Natl Acad Sci U S A 106, 4707–4712.

Bianchini, G., De Angelis, C., Licata, L., and Gianni, L. (2022). Treatment landscape of triple-negative breast cancer - expanded options, evolving needs. Nat Rev Clin Oncol 19, 91–113.

Breton, B., Sauvageau, E., Zhou, J., Bonin, H., Le Gouill, C., and Bouvier, M. (2010). Multiplexing of multicolor bioluminescence resonance energy transfer. Biophys J 99, 4037–4046.

Bruce, B., Khanna, G., Ren, L., Landberg, G., Jirstrom, K., Powell, C., Borczuk, A., Keller, E.T., Wojno, K.J., Meltzer, P., et al. (2007). Expression of the cytoskeleton linker protein ezrin in human cancers. Clinical & experimental metastasis 24, 69–78.

Cant, S.H., and Pitcher, J.A. (2005). G protein-coupled receptor kinase 2-mediated phosphorylation of ezrin is required for G protein-coupled receptor-dependent reorganization of the actin cytoskeleton. Mol Biol Cell 16, 3088–3099.

Carreno, S., Kouranti, I., Glusman, E.S., Fuller, M.T., Echard, A., and Payre, F. (2008). Moesin and its activating kinase Slik are required for cortical stability and microtubule organization in mitotic cells. J Cell Biol 180, 739–746.

Clark, A.G., and Vignjevic, D.M. (2015). Modes of cancer cell invasion and the role of the microenvironment. Curr Opin Cell Biol 36, 13–22.

Clucas, J., and Valderrama, F. (2014). ERM proteins in cancer progression. J Cell Sci 127, 267–275.

Coleman, R.A., Humphrey, P.P., Kennedy, I., Levy, G.P., and Lumley, P. (1981). Comparison of the actions of U-46619, a prostaglandin H2-analogue, with those of prostaglandin H2 and thromboxane A2 on some isolated smooth muscle preparations. British journal of pharmacology 73, 773–778.

Curto, M., and McClatchey, A.I. (2004). Ezrin…a metastatic detERMinant? Cancer Cell 5, 113–114.

De Jamblinne, C.V., Decelle, B., Dehghani, M., Joseph, M., Sriskandarajah, N., Leguay, K., Rambaud, B., Lemieux, S., Roux, P.P., Hipfner, D.R., et al. (2020). STRIPAK regulates Slik localization to control mitotic morphogenesis and epithelial integrity. J Cell Biol 219.

Devost, D., Sleno, R., Petrin, D., Zhang, A., Shinjo, Y., Okde, R., Aoki, J., Inoue, A., and Hebert, T.E. (2017). Conformational Profiling of the AT1 Angiotensin II Receptor Reflects Biased Agonism, G Protein Coupling, and Cellular Context. J Biol Chem 292, 5443–5456.

Ekambaram, P., Lambiv, W., Cazzolli, R., Ashton, A.W., and Honn, K.V. (2011). The thromboxane synthase and receptor signaling pathway in cancer: an emerging paradigm in cancer progression and metastasis. Cancer metastasis reviews 30, 397–408.

Elliott, B.E., Meens, J.A., SenGupta, S.K., Louvard, D., and Arpin, M. (2005). The membrane cytoskeletal crosslinker ezrin is required for metastasis of breast carcinoma cells. Breast cancer research : BCR 7, R365–373.

Estecha, A., Sanchez-Martin, L., Puig-Kroger, A., Bartolome, R.A., Teixido, J., Samaniego, R., and Sanchez-Mateos, P. (2009). Moesin orchestrates cortical polarity of melanoma tumour cells to initiate 3D invasion. J Cell Sci 122, 3492–3501.

Fehon, R.G., McClatchey, A.I., and Bretscher, A. (2010). Organizing the cell cortex: the role of ERM proteins. Nat Rev Mol Cell Biol 11, 276–287.

Fife, C.M., McCarroll, J.A., and Kavallaris, M. (2014). Movers and shakers: cell cytoskeleton in cancer metastasis. British journal of pharmacology 171, 5507–5523.

Gales, C., Rebois, R.V., Hogue, M., Trieu, P., Breit, A., Hebert, T.E., and Bouvier, M. (2005). Real-time monitoring of receptor and G-protein interactions in living cells. Nat Methods 2, 177–184.

Gary, R., and Bretscher, A. (1995). Ezrin self-association involves binding of an N-terminal domain to a normally masked C-terminal domain that includes the F-actin binding site. Mol Biol Cell 6, 1061–1075.

Ghaffari, A., Hoskin, V., Turashvili, G., Varma, S., Mewburn, J., Mullins, G., Greer, P.A., Kiefer, F., Day, A.G., Madarnas, Y., et al. (2019). Intravital imaging reveals systemic ezrin inhibition impedes cancer cell migration and lymph node metastasis in breast cancer. Breast cancer research : BCR 21, 12.

Guillem-Llobat, P., Dovizio, M., Bruno, A., Ricciotti, E., Cufino, V., Sacco, A., Grande, R., Alberti, S., Arena, V., Cirillo, M., et al. (2016). Aspirin prevents colorectal cancer metastasis in mice by splitting the crosstalk between platelets and tumor cells. Oncotarget 7, 32462–32477.

Hoskin, V., Ghaffari, A., and Elliott, B.E. (2019). Ezrin, more than a metastatic detERMinant? Oncotarget 10, 6755–6757.

Hoskin, V., Szeto, A., Ghaffari, A., Greer, P.A., Cote, G.P., and Elliott, B.E. (2015). Ezrin regulates focal adhesion and invadopodia dynamics by altering calpain activity to promote breast cancer cell invasion. Mol Biol Cell 26, 3464–3479.

Jones, R.L., Wilson, N.H., and Armstrong, R.A. (1985). Characterization of thromboxane receptors in human platelets. Adv Exp Med Biol 192, 67–81.

Kattelman, E.J., Venton, D.L., and Le Breton, G.C. (1986). Characterization of U46619 binding in unactivated, intact human platelets and determination of binding site affinities of four TXA2/PGH2 receptor antagonists (13-APA, BM 13.177, ONO 3708 and SQ 29,548). Thrombosis research 41, 471–481.

Khanna, C., Wan, X., Bose, S., Cassaday, R., Olomu, O., Mendoza, A., Yeung, C., Gorlick, R., Hewitt, S.M., and Helman, L.J. (2004). The membrane-cytoskeleton linker ezrin is necessary for osteosarcoma metastasis. Nat Med 10, 182–186.

Kobayashi, H., Picard, L.P., Schonegge, A.M., and Bouvier, M. (2019). Bioluminescence resonance energy transfer-based imaging of protein-protein interactions in living cells. Nature protocols 14, 1084–1107.

Kobayashi, H., Sagara, J., Kurita, H., Morifuji, M., Ohishi, M., Kurashina, K., and Taniguchi, S. (2004). Clinical significance of cellular distribution of moesin in patients with oral squamous cell carcinoma. Clinical Cancer Research 10, 572–580.

Koedoot, E., Fokkelman, M., Rogkoti, V.M., Smid, M., van de Sandt, I., de Bont, H., Pont, C., Klip, J.E., Wink, S., Timmermans, M.A., et al. (2019). Uncovering the signaling landscape controlling breast cancer cell migration identifies novel metastasis driver genes. Nature communications 10, 2983.

Kunda, P., Pelling, A.E., Liu, T., and Baum, B. (2008). Moesin controls cortical rigidity, cell rounding, and spindle morphogenesis during mitosis. Curr Biol 18, 91–101.

Leguay, K., Decelle, B., Elkholi, I.E., Bouvier, M., Cote, J.F., and Carreno, S. (2022). Interphase microtubule disassembly is a signaling cue that drives cell rounding at mitotic entry. J Cell Biol 221.

Leguay, K., Decelle, B., He, Y.Y., Pagniez, A., Hogue, M., Kobayashi, H., Le Gouill, C., Bouvier, M., and Carreno, S. (2021). Development of conformational BRET biosensors that monitor ezrin, radixin and moesin activation in real time. J Cell Sci 134.

Li, H., Lee, M.H., Liu, K., Wang, T., Song, M., Han, Y., Yao, K., Xie, H., Zhu, F., Grossmann, M., et al. (2017). Inhibiting breast cancer by targeting the thromboxane A2 pathway. NPJ Precis Oncol 1, 8.

Liu, D., Ge, L., Wang, F., Takahashi, H., Wang, D., Guo, Z., Yoshimura, S.H., Ward, T., Ding, X., Takeyasu, K., et al. (2007). Single-molecule detection of phosphorylation-induced plasticity changes during ezrin activation. FEBS Lett 581, 3563–3571.

Lucotti, S., Cerutti, C., Soyer, M., Gil-Bernabe, A.M., Gomes, A.L., Allen, P.D., Smart, S., Markelc, B., Watson, K., Armstrong, P.C., et al. (2019). Aspirin blocks formation of metastatic intravascular niches by inhibiting platelet-derived COX-1/thromboxane A2. J Clin Invest 129, 1845–1862.

Lukasheva, V., Devost, D., Le Gouill, C., Namkung, Y., Martin, R.D., Longpre, J.M., Amraei, M., Shinjo, Y., Hogue, M., Lagace, M., et al. (2020). Signal profiling of the beta1AR reveals coupling to novel signalling pathways and distinct phenotypic responses mediated by beta1AR and beta2AR. Sci Rep 10, 8779.

Lutz, S., Shankaranarayanan, A., Coco, C., Ridilla, M., Nance, M.R., Vettel, C., Baltus, D., Evelyn, C.R., Neubig, R.R., Wieland, T., et al. (2007). Structure of Galphaq- p63RhoGEF-RhoA complex reveals a pathway for the activation of RhoA by GPCRs. Science 318, 1923–1927.

Machicoane, M., de Frutos, C.A., Fink, J., Rocancourt, M., Lombardi, Y., Garel, S., Piel, M., and Echard, A. (2014). SLK-dependent activation of ERMs controls LGN-NuMA localization and spindle orientation. J Cell Biol 205, 791–799.

Matsui, T., Amano, M., Yamamoto, T., Chihara, K., Nakafuku, M., Ito, M., Nakano, T., Okawa, K., Iwamatsu, A., and Kaibuchi, K. (1996). Rho-associated kinase, a novel serine/threonine kinase, as a putative target for small GTP binding protein Rho. EMBO J 15, 2208–2216.

Matsui, T., Maeda, M., Doi, Y., Yonemura, S., Amano, M., Kaibuchi, K., Tsukita, S., and Tsukita, S. (1998). Rho-kinase phosphorylates COOH-terminal threonines of ezrin/radixin/moesin (ERM) proteins and regulates their head-to-tail association. J Cell Biol 140, 647–657.

Meiri, D., Marshall, C.B., Mokady, D., LaRose, J., Mullin, M., Gingras, A.C., Ikura, M., and Rottapel, R. (2014). Mechanistic insight into GPCR-mediated activation of the microtubule-associated RhoA exchange factor GEF-H1. Nature communications 5, 4857.

Nakahata, N. (2008). Thromboxane A2: physiology/pathophysiology, cellular signal transduction and pharmacology. Pharmacology & therapeutics 118, 18–35.

Nakamura, F., Amieva, M.R., and Furthmayr, H. (1995). Phosphorylation of threonine 558 in the carboxyl-terminal actin-binding domain of moesin by thrombin activation of human platelets. J Biol Chem 270, 31377–31385.

Namkung, Y., Le Gouill, C., Lukashova, V., Kobayashi, H., Hogue, M., Khoury, E., Song, M., Bouvier, M., and Laporte, S.A. (2016). Monitoring G protein-coupled receptor and beta-arrestin trafficking in live cells using enhanced bystander BRET. Nature communications 7, 12178.

Namkung, Y., LeGouill, C., Kumar, S., Cao, Y., Teixeira, L.B., Lukasheva, V., Giubilaro, J., Simoes, S.C., Longpre, J.M., Devost, D., et al. (2018). Functional selectivity profiling of the angiotensin II type 1 receptor using pathway-wide BRET signaling sensors. Science signaling 11.

Nie, D., Che, M., Zacharek, A., Qiao, Y., Li, L., Li, X., Lamberti, M., Tang, K., Cai, Y., Guo, Y., et al. (2004). Differential expression of thromboxane synthase in prostate carcinoma: role in tumor cell motility. Am J Pathol 164, 429–439.

Ogletree, M.L., Harris, D.N., Greenberg, R., Haslanger, M.F., and Nakane, M. (1985). Pharmacological actions of SQ 29,548, a novel selective thromboxane antagonist. The Journal of pharmacology and experimental therapeutics 234, 435–441.

Orr, K., Buckley, N.E., Haddock, P., James, C., Parent, J.L., McQuaid, S., and Mullan, P.B. (2016). Thromboxane A2 receptor (TBXA2R) is a potent survival factor for triple negative breast cancers (TNBCs). Oncotarget 7, 55458–55472.

Pang, S.T., Fang, X., Valdman, A., Norstedt, G., Pousette, A., Egevad, L., and Ekman, P. (2004). Expression of ezrin in prostatic intraepithelial neoplasia. Urology 63, 609–612.

Papatheodorou, I., Moreno, P., Manning, J., Fuentes, A.M., George, N., Fexova, S., Fonseca, N.A., Fullgrabe, A., Green, M., Huang, N., et al. (2020). Expression Atlas update: from tissues to single cells. Nucleic Acids Res 48, D77–D83.

Parent, J.L., Labrecque, P., Orsini, M.J., and Benovic, J.L. (1999). Internalization of the TXA2 receptor alpha and beta isoforms. Role of the differentially spliced cooh terminus in agonist-promoted receptor internalization. J Biol Chem 274, 8941–8948.

Park, J., Jang, J.H., Oh, S., Kim, M., Shin, C., Jeong, M., Heo, K., Park, J.B., Kim, S.R., and Oh, Y.S. (2018). LPA-induced migration of ovarian cancer cells requires activation of ERM proteins via LPA1 and LPA2. Cellular signalling 44, 138–147.

Pulix, M., Lukashchuk, V., Smith, D.C., and Dickson, A.J. (2021). Molecular characterization of HEK293 cells as emerging versatile cell factories. Current opinion in biotechnology 71, 18–24.

Qin, Y., Chen, W., Jiang, G., Zhou, L., Yang, X., Li, H., He, X., Wang, H.L., Zhou, Y.B., Huang, S., et al. (2020). Interfering MSN-NONO complex-activated CREB signaling serves as a therapeutic strategy for triple-negative breast cancer. Sci Adv 6, eaaw9960.

Ren, L., and Khanna, C. (2014). Role of ezrin in osteosarcoma metastasis. Adv Exp Med Biol 804, 181–201.

Retzer, M., and Essler, M. (2000). Lysophosphatidic acid-induced platelet shape change proceeds via Rho/Rho kinase-mediated myosin light-chain and moesin phosphorylation. Cellular signalling 12, 645–648.

Roubinet, C., Decelle, B., Chicanne, G., Dorn, J.F., Payrastre, B., Payre, F., and Carreno, S. (2011). Molecular networks linked by Moesin drive remodeling of the cell cortex during mitosis. J Cell Biol 195, 99–112.

Sah, V.P., Seasholtz, T.M., Sagi, S.A., and Brown, J.H. (2000). The role of Rho in G protein-coupled receptor signal transduction. Annual review of pharmacology and toxicology 40, 459–489.

Schrage, R., Schmitz, A.L., Gaffal, E., Annala, S., Kehraus, S., Wenzel, D., Bullesbach, K.M., Bald, T., Inoue, A., Shinjo, Y., et al. (2015). The experimental power of FR900359 to study Gq-regulated biological processes. Nature communications 6, 10156.

Seyfried, T.N., and Huysentruyt, L.C. (2013). On the origin of cancer metastasis. Critical reviews in oncogenesis 18, 43–73.

Shaw, R.J., Henry, M., Solomon, F., and Jacks, T. (1998). RhoA-dependent phosphorylation and relocalization of ERM proteins into apical membrane/actin protrusions in fibroblasts. Mol Biol Cell 9, 403–419.

Shcherbina, A., Kenney, D.M., Bretscher, A., and Remold, O.D.E. (1999). Dynamic association of moesin with the membrane skeleton of thrombin- activated platelets. Blood 93, 2128–2129.

Simons, P.C., Pietromonaco, S.F., Reczek, D., Bretscher, A., and Elias, L. (1998). C-terminal threonine phosphorylation activates ERM proteins to link the cell’s cortical lipid bilayer to the cytoskeleton. Biochem Biophys Res Commun 253, 561–565.

Solinet, S., Mahmud, K., Stewman, S.F., Ben El Kadhi, K., Decelle, B., Talje, L., Ma, A., Kwok, B.H., and Carreno, S. (2013). The actin-binding ERM protein Moesin binds to and stabilizes microtubules at the cell cortex. J Cell Biol 202, 251–260.

Song, Y., Ma, X., Zhang, M., Wang, M., Wang, G., Ye, Y., and Xia, W. (2020). Ezrin Mediates Invasion and Metastasis in Tumorigenesis: A Review. Front Cell Dev Biol 8, 588801.

Takasaki, J., Saito, T., Taniguchi, M., Kawasaki, T., Moritani, Y., Hayashi, K., and Kobori, M. (2004). A novel Galphaq/11-selective inhibitor. J Biol Chem 279, 47438–47445.

Tsherniak, A., Vazquez, F., Montgomery, P.G., Weir, B.A., Kryukov, G., Cowley, G.S., Gill, S., Harrington, W.F., Pantel, S., Krill-Burger, J.M., et al. (2017). Defining a Cancer Dependency Map. Cell 170, 564–576 e516.

Vaiskunaite, R., Adarichev, V., Furthmayr, H., Kozasa, T., Gudkov, A., and Voyno-Yasenetskaya, T.A. (2000). Conformational activation of radixin by G13 protein alpha subunit. J Biol Chem 275, 26206–26212.

van Unen, J., Yin, T., Wu, Y.I., Mastop, M., Gadella, T.W., Jr., and Goedhart, J. (2016). Kinetics of recruitment and allosteric activation of ARHGEF25 isoforms by the heterotrimeric G-protein Galphaq. Sci Rep 6, 36825.

Viswanatha, R., Ohouo, P.Y., Smolka, M.B., and Bretscher, A. (2012). Local phosphocycling mediated by LOK/SLK restricts ezrin function to the apical aspect of epithelial cells. J Cell Biol 199, 969–984.

Watkins, G., Douglas-Jones, A., Mansel, R.E., and Jiang, W.G. (2005). Expression of thromboxane synthase, TBXAS1 and the thromboxane A2 receptor, TBXA2R, in human breast cancer. Int Semin Surg Oncol 2, 23.

Welch, D.R., and Hurst, D.R. (2019). Defining the Hallmarks of Metastasis. Cancer Res 79, 3011–3027.

Werfel, T.A., Hicks, D.J., Rahman, B., Bendeman, W.E., Duvernay, M.T., Maeng, J.G., Hamm, H., Lavieri, R.R., Joly, M.M., Pulley, J.M., et al. (2020). Repurposing of a Thromboxane Receptor Inhibitor Based on a Novel Role in Metastasis Identified by Phenome-Wide Association Study. Molecular cancer therapeutics 19, 2454–2464.

Wikstrom, K., Kavanagh, D.J., Reid, H.M., and Kinsella, B.T. (2008). Differential regulation of RhoA-mediated signaling by the TPalpha and TPbeta isoforms of the human thromboxane A2 receptor: independent modulation of TPalpha signaling by prostacyclin and nitric oxide. Cellular signalling 20, 1497–1512.

Wilde, C., and Aktories, K. (2001). The Rho-ADP-ribosylating C3 exoenzyme from Clostridium botulinum and related C3-like transferases. Toxicon 39, 1647–1660.

Yankaskas, C.L., Thompson, K.N., Paul, C.D., Vitolo, M.I., Mistriotis, P., Mahendra, A., Bajpai, V.K., Shea, D.J., Manto, K.M., Chai, A.C., et al. (2019). A microfluidic assay for the quantification of the metastatic propensity of breast cancer specimens. Nat Biomed Eng 3, 452–465.

Yousef, E.M., Tahir, M.R., St-Pierre, Y., and Gaboury, L.A. (2014). MMP-9 expression varies according to molecular subtypes of breast cancer. BMC cancer 14, 609.

Yu, O.M., and Brown, J.H. (2015). G Protein-Coupled Receptor and RhoA-Stimulated Transcriptional Responses: Links to Inflammation, Differentiation, and Cell Proliferation. Molecular pharmacology 88, 171–180.

Yu, Y., Khan, J., Khanna, C., Helman, L., Meltzer, P.S., and Merlino, G. (2004). Expression profiling identifies the cytoskeletal organizer ezrin and the developmental homeoprotein Six-1 as key metastatic regulators. Nat Med 10, 175–181.

